# Root hair-endophyte stacking (RHESt) in an ancient Afro-Indian crop creates an unusual physico-chemical barrier to trap pathogen(s)

**DOI:** 10.1101/071548

**Authors:** W. K. Mousa, C. Shearer, Victor Limay-Rios, C. Ettinger, J. A. Eisen, M.N. Raizada

**Affiliations:** Department of Plant Agriculture, University of Guelph, Guelph, ON Canada N1G 2W1; Department of Plant Agriculture, University of Guelph, Ridgetown Campus, Ridgetown, ON, 7 Canada, N0P 2C0; University of California Davis Genome Center, Davis, California, USA 95616

## Abstract

The ancient African crop, finger millet, has broad resistance to pathogens including the toxigenic fungus *Fusarium graminearum*. Here we report the discovery of a novel plant defence mechanism, resulting from an unusual symbiosis between finger millet and a root-inhabiting bacterial endophyte, M6 (*Enterobacter* sp.). Seed-coated M6 swarms towards *Fusarium* attempting to penetrate root epidermis, induces growth of root hairs which then bend parallel to the root axis, then forms biofilm-mediated microcolonies, resulting in a remarkable, multi-layer root hair-endophyte stack (RHESt). RHESt results in a physical barrier that prevents entry and/or traps *F. graminearum* which is then killed. Thus M6 creates its own specialized killing microhabitat. M6 killing requires c-di-GMP-dependent signalling, diverse fungicides and xenobiotic resistance. Further molecular evidence suggests long-term host-endophyte-pathogen co-evolution. The end-result of this remarkable symbiosis is reduced DON mycotoxin, potentially benefiting millions of subsistence farmers and livestock. RHESt demonstrates the value of exploring ancient, orphan crop microbiomes.

## Introduction

Finger millet (*Eleusine coracana*) is a cereal crop widely grown by subsistence farmers in 36 Africa and South Asia ^1,2^. Finger millet was domesticated in Western Uganda and Ethiopia around 5000 BC then reached India by 3000 BC ^3^. With few exceptions, subsistence farmers report that finger millet is widely resistant to pathogens including *Fusarium* species ^4,5^. One species of *Fusarium*, *F. graminearum,* causes devastating diseases in crops related to finger millet including maize, wheat and barley, associated with accumulation of the mycotoxin deoxynivalenol (DON) which affects humans and livestock ^6,7^. However, despite its prevalence as a disease-causing agent across cereals, *F. graminearum* is not considered to be an important pathogen of finger millet, suggesting this crop has tolerance to this *Fusarium* species ^4,8^.

The resistance of finger millet grain to fungal disease has been attributed to high concentrations of polyphenols ^9,10^. However, emerging literature suggests that microbes that inhabit plants without themselves causing disease, defined as endophytes, may contribute to host resistance against fungal pathogens ^11,12^. Endophytes have been shown to suppress fungal diseases through induction of host resistance genes ^13^, competition ^14^, and/or production of anti-pathogenic natural compounds ^15,16^.

*Fusarium* are ancient fungal species, dating to at least 8.8 million years ago, and their diversification appears to have co-occurred with that of the C4 grasses (which includes finger millet), certainly pre-dating finger millet domestication in Africa ^17^. Multiple studies have reported the presence of *Fusarium* in finger millet in Africa and India^18-23^. A diversity of *F. verticillioides* (synonym *F. moniliforme*) has been observed in finger millet in Africa and it has been suggested that the species evolved there ^18^. These observations suggest the possibility of co-evolution within finger millet between *Fusarium* and millet endophytes. We previously isolated, for the first time, fungal endophytes from finger millet and showed that their natural products have anti-*Fusarium* activity ^4^. We could not find reports of bacterial endophytes isolated from 61 finger millet.

The objectives of this study were to isolate bacterial endophytes from finger millet, assay for anti-*Fusarium* activity and characterize the underlying cellular, molecular and biochemical mechanisms. We report an unusual symbiosis between the host and a root-inhabiting bacterial endophyte.

## Results

### Isolation, identification and antifungal activity of endophytes

A total of seven bacterial endophytes were isolated from surface-sterilized tissues of finger millet, strains M1 to M7 (Fig. 1a-c and Supplementary Table 1). BLAST searching of the 16S rDNA sequences against Genbank suggested that strains M1 and M6 resemble *Enterobacter* sp. while M2 and M4 resemble *Pantoea* sp.., and M3, M5, and M7 resemble *Burkholderia* sp..GenBank accession numbers for strains M1, M2, M3, M4, M5, M6 and M7 are KU307449, KU307450, KU307451, KU307452, KU307453, KU307454, and KU307455, respectively. The 16S rDNA sequences for finger millet bacterial endophytes were used to generate a phylogenetic tree (Supplementary Fig. 1) using Phylogeny.fr software ^24,25^. Interestingly, five of the seven strains showed anti-fungal activity against *F. graminearum in vitro* (Fig. 1c-d). Strain M6 from millet roots showing the most potent activity and hence was selected for further study including whole genome sequencing, which resulted in a final taxonomic classification (*Enterobacter* sp., strain UCD-UG_FMILLET) ^26^. M6 was observed to inhibit the growth of 5 out of 20 additional crop-associated fungi, including pathogens, suggesting it has a wider target spectrum (Supplementary Table 2). As viewed by electron microscopy, M6 showed an elongated rod shape with a wrinkled surface (Fig. 1e). Following seed coating, GFP-tagged M6 localized to finger millet roots intercellularly and intracellularly (Fig. 1f,g). In addition, colonization of finger millet with M6 did not result in pathogenic symptoms (Supplementary Fig. 2). Combined, these results confirm that M6 is an endophyte of finger millet.

**Figure 1.**
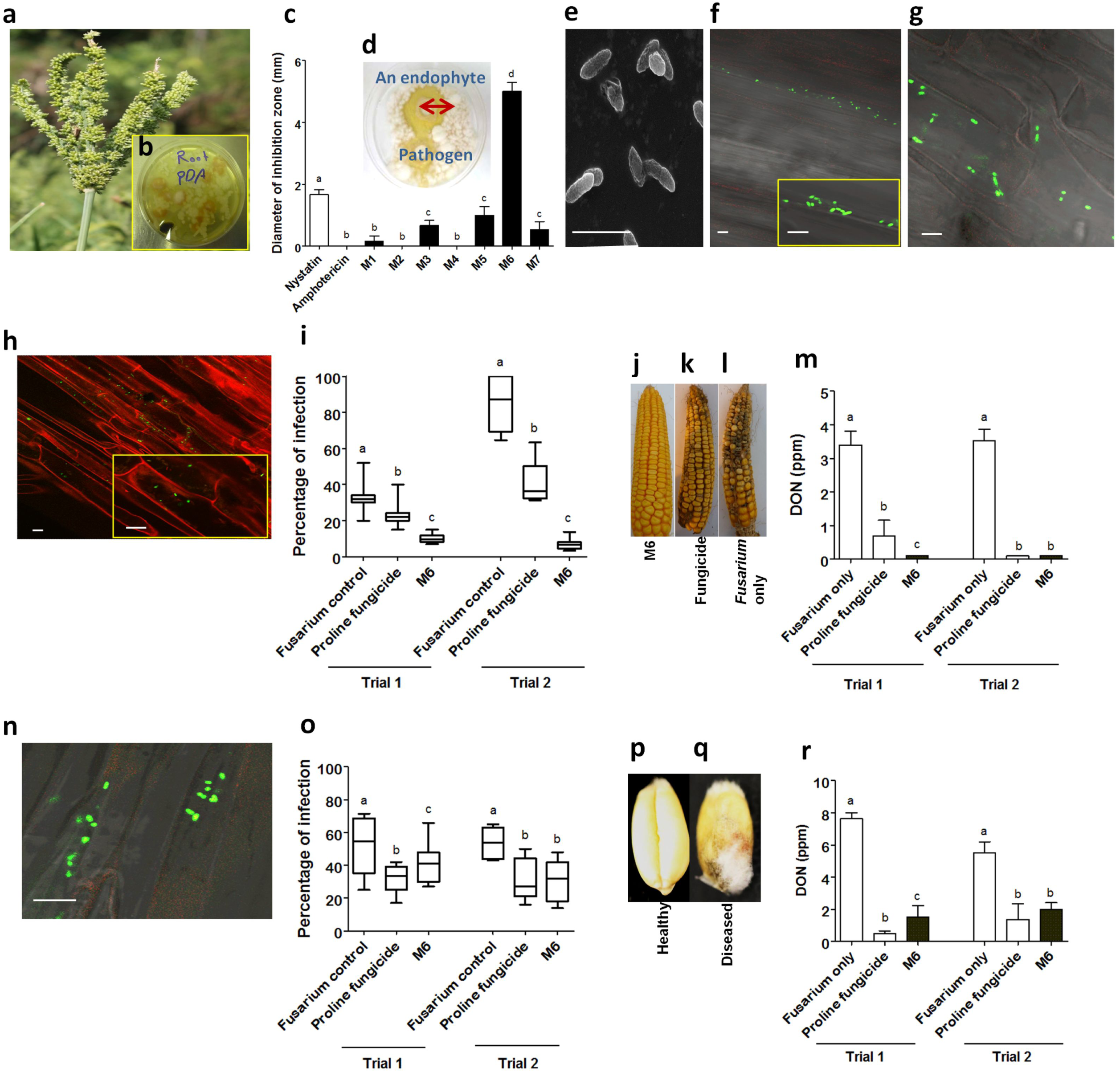
Isolation, identification and antifungal activity of endophytes. **a**, Picture showing finger millet grain head. **b**, Mixed culture of endophytes isolated from finger millet. **c**, Quantification of the inhibitory effect of finger millet endophytes or fungicide controls on the growth of *F. graminearum in vitro.* For these experiments, n=3. **d**, M6 endophyte suppresses the growth of *F. graminearum* hyphae (white) using the dual culture method. **e**, Imaging of M6 viewed by scanning electron microscopy. **f-g**, GFP-tagged M6 inside roots of finger millet viewed by scanning confocal microscopy. **h**, GFP-tagged M6 inside roots of maize (stained with propidium iodide). **i**, Effect of M6 treatment on suppression of *F. graminearum* disease in maize in two greenhouse trials. **j-l**, (left to right) Representative ears from M6, fungicide and *Fusarium* only treatments. **m**, Effect of M6 or controls on reducing DON mycotoxin contamination in maize during storage following the two greenhouse trials. **n**, GFP-tagged M6 inside roots of wheat viewed by confocal microscopy. **o**, Effect of M6 treatment on suppression of *F. graminearum* disease in wheat in two greenhouse trials. **p**, Picture of a healthy wheat grain. **q**, Picture of an infected wheat grain. **r**, Effect of M6 or controls on reducing DON mycotoxin contamination in wheat during storage following the two greenhouse trials. Scale bars in all pictures equal 5 µm. For greenhouse disease trials, n=20 for M6 and n=10 for the controls. For DON quantification, n=3 pools of seeds. The whiskers (**i, o**) indicate the range of data points. The error bars (**c, m, q**) indicate the standard error of the mean. For all graphs, letters that are different from one another indicate that their means are statistically different (P≤0.05).

**Figure 2.**
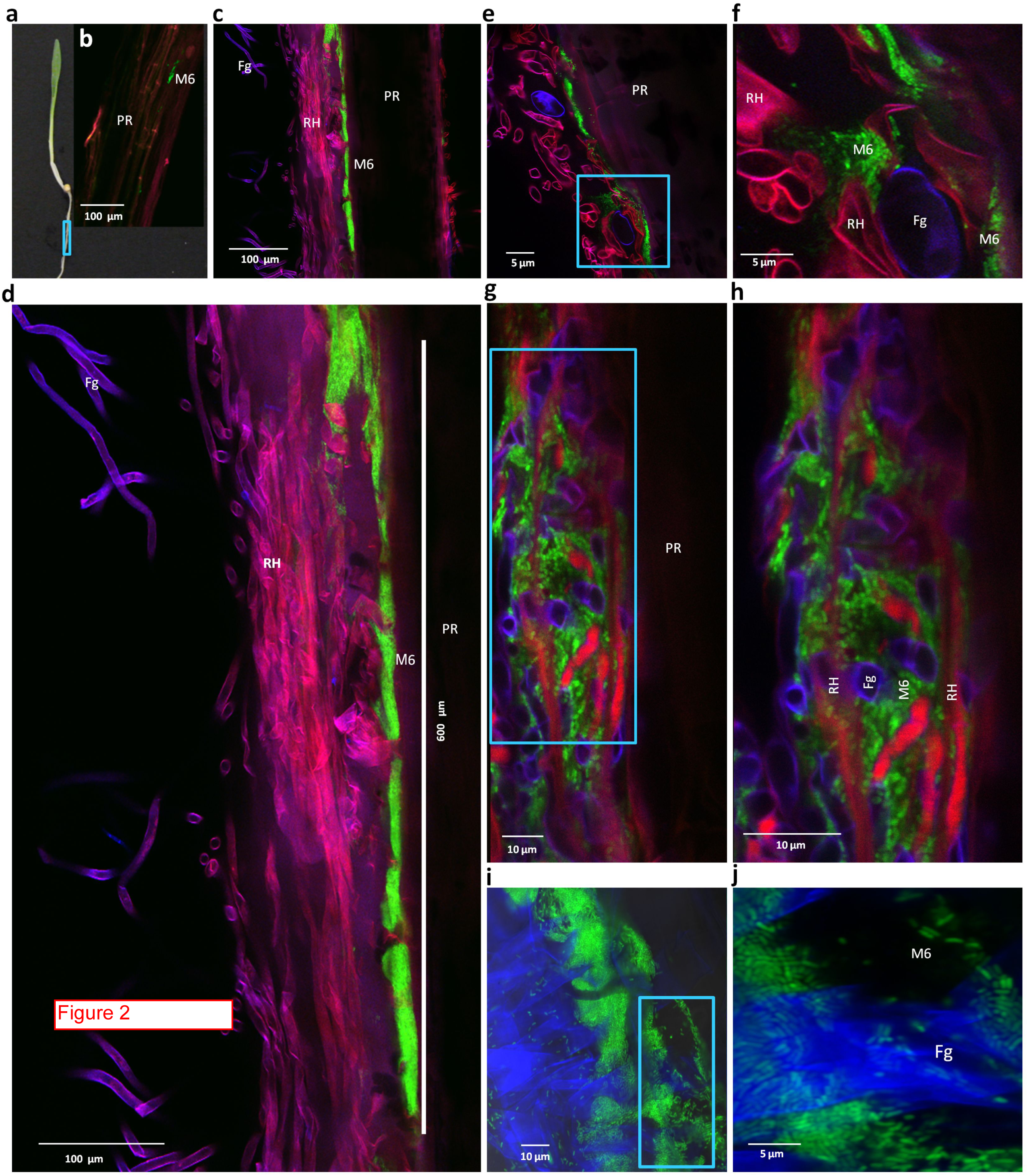
Confocal imaging of m6-*fusarium* interactions in finger millet roots. **a**, Picture of millet seedling showing primary root (PR) zone used for confocal microscopy. **b**, Control primary root that was seed coated with GFP-M6 (green) but not infected with *F. graminearum*. As a control, the tissue was stained with fungal stain calcofluor to exclude the presence of other fungi. Root following seed coating with GFP-tagged endophyte M6 (M6, green) following inoculation with *F. graminearum* (Fg, purple blue, calcofluor stained) showing interactions with root hairs (RH, magenta, lignin autofluoresence). **c**-**d,** Low (**c**) and high (**d**) magnifications to show the dense root hair and endophyte barrier layers. **e**-**f**, Low (**e**) and high (**f**) magnifications at the edge of the barrier layers. **g-h**, Low (**g**) and high (**h**) magnifications in a deeper confocal plane of the root hair layer shown in (**d**) showing root hair endophyte stacking (RHESt) with trapped fungal hyphae. **i-j**, Low (**i**) and high (**j**) magnifications of the interactions between M6 (green) and *F. graminearum* in the absence of root hair-lignin autofluorescence, showing breakage of fungal hyphae.

To determine whether strain M6 has anti-*Fusarium* activity *in planta*, related *Fusarium*-susceptible cereals (maize and wheat) were used as model systems (Fig. 1 h-r), since finger millet itself is not reported to be susceptible to *F. graminearum.* Seed-coated GFP-tagged M6 was shown to colonize the internal tissues of maize (Fig. 1h) and wheat (Fig. 1n) suggesting it can also behave as an endophyte in these crop relatives. Treatments (combined seed coating and foliar spray) with M6 caused statistically significant (P≤0.05) reductions in *F. graminearum* disease symptoms in maize (Gibberella Ear Rot, Fig. 1i-l) and wheat (Fusarium Head Blight, Fig. 1o-q) ranging from 70 to 90% and 20-30%, respectively in two greenhouse trials, compared to plants treated with *Fusarium* only (yield data in Supplementary Fig. 3a-b; Supplementary Table 3). Foliar spraying alone with M6 resulted in more disease reduction compared to seed coating alone, though this effect was not statistically significant, at P≤0.05 (Supplementary Fig. 3c). 100 Following extended storage to mimic those of African subsistence farmers (ambient temperature and moisture), treatment with M6 resulted in dramatic reductions in DON accumulation, with DON levels declining from ∼3.4 ppm to 0.1 ppm in maize, and from 5.5 ppm to 0.2 ppm in wheat, equivalent to 97% and 60% reductions, respectively, compared to plants treated with *Fusarium* only, at P≤0.05 (Fig. 1m,r and Supplementary Table 4).

**Figure 3.**
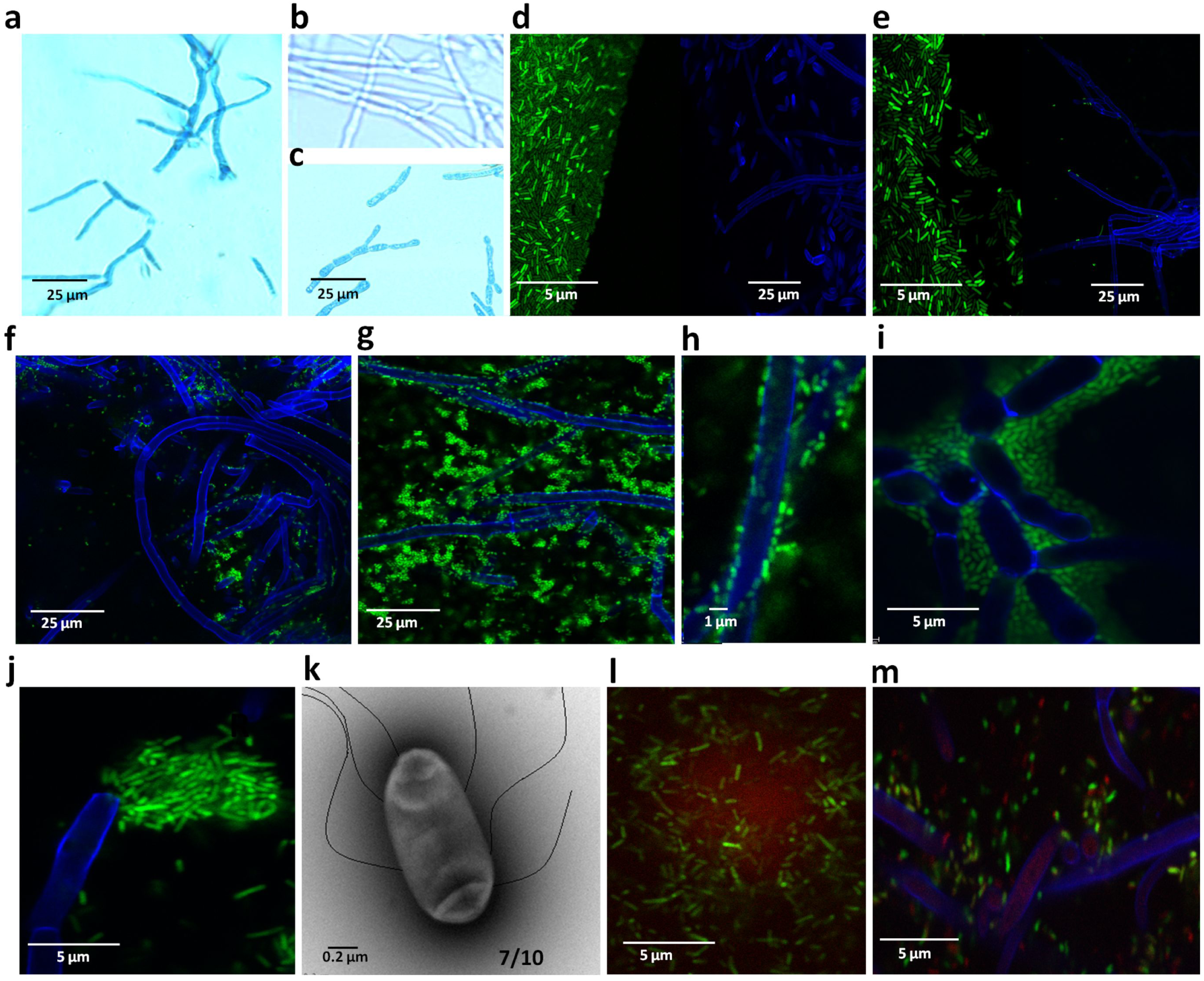
Behavior and interactions of endophyte m6 and *f. graminearum in vitro* on microscope slides. **a-c**, Light microscopy of interactions between *F. graminearum* (Fg) and M6 following staining with Evans blue, which stains dead hyphae blue. Shown are (**a**) Fg following overnight co-incubation with M6, (**b**) Fg, grown away from M6 (control), and (**c**) Fg following overnight co-incubation with a commercial biological control agent. **d-i**, Time course to illustrate the swarming and attachment behaviour of GFP-tagged M6 (green) to Fg (blue, calcofluor stained) viewed at 0.5 h, 1.5 h, 3 h, 6 h, 6 h (close-up) and 8 h following co-incubation, respectively. Fg and M6 shown in (**d**) and (**e**) were inoculated on the same slide distal from one another at the start of the time course but digitally placed together for these illustrations. **j,** M6 shown breaking Fg hypha. **k,** Transmission electron microscope picture of M6 showing its characteristic flagella. (**l-m**) Biofilm formed by M6 as viewed by staining with Ruby film tracer (red) in the (**l**) absence of Fg or (**m**) presence of Fg.

### Microscopic imaging of M6-*Fusarium* interactions in finger millet roots and *in vitro*

Since *F. graminearum* has been reported to infect cereal roots ^27^, and since endophyte M6 was originally isolated from the same tissue, finger millet roots were selected to visualize potential interactions between M6 and *F. graminearum* (Fig. 2). GFP-tagged M6 was coated onto millet seed. Following germination (Fig. 2a), GFP-M6 showed sporadic, low population density distribution throughout the seminal roots (Fig. 2b). Following inoculation with *F. graminearum,* which was visualized using calcofluor staining, GFP-M6 accumulated at sites of attempted entry by *Fusarium,* creating a remarkable, high density layer of microcolonies of M6 along the entire root epidermal surface, the rhizoplane (Fig. 2c,d). External to the M6 rhizoplane barrier was a thick mat of root hairs (RH) (Fig. 2c,d). RH number and length were much greater at sites of M6 accumulation compared to the opposite side (Fig. 2c), and M6 was shown *in vitro* to produce auxin (Supplemental Method 1, Supplementary Fig. 4), a known RH-growth promoting plant hormone ^28^. Interestingly, most RH were bent, parallel to the root axis (Fig. 2d). The RH mat appeared to obscure M6 cells, and when observed in a low RH density area (Fig. 2e), M6 cells were clearly visible and appeared to attach onto root hairs and engulf *Fusarium* hyphae (Fig. 2f). Imaging at deeper confocal planes below the surface of the RH mat (Fig. 2g,h) revealed that the mat did not consist only of RH, but rather that M6 cells were intercalated between bent RH strands forming an unusual, multi-layer root hair-endophyte stack (RHESt). Within the RHESt, *F. graminearum* hyphae appeared to be trapped (Fig. 2h). By imaging only the M6-*Fusarium* interaction within the RHESt, M6 microcolonies were observed to be associated with breakage of the fungal hyphae (Fig. 2i,j). To confirm that the endophyte actively kills *Fusarium,* Evans blue vitality stain, which stains dead hyphae blue, was used following co-incubation on a microscope slide. The fungal hyphae in contact with strain M6 stained blue and appeared broken (Fig. 3a) in contrast to the control (*F. graminearum* exposed to buffer only) (Fig. 3b). The M6 result was similar to the well known fungicidal activity of the commercial biocontrol agent, *B. subtilis* (Fig. 3c). Combined, these results suggest that M6 cooperates with RH cells to create a specialized killing microhabitat (RHESt) that protects millet roots from invasion by *F. graminearum.*

**Figure 4.**
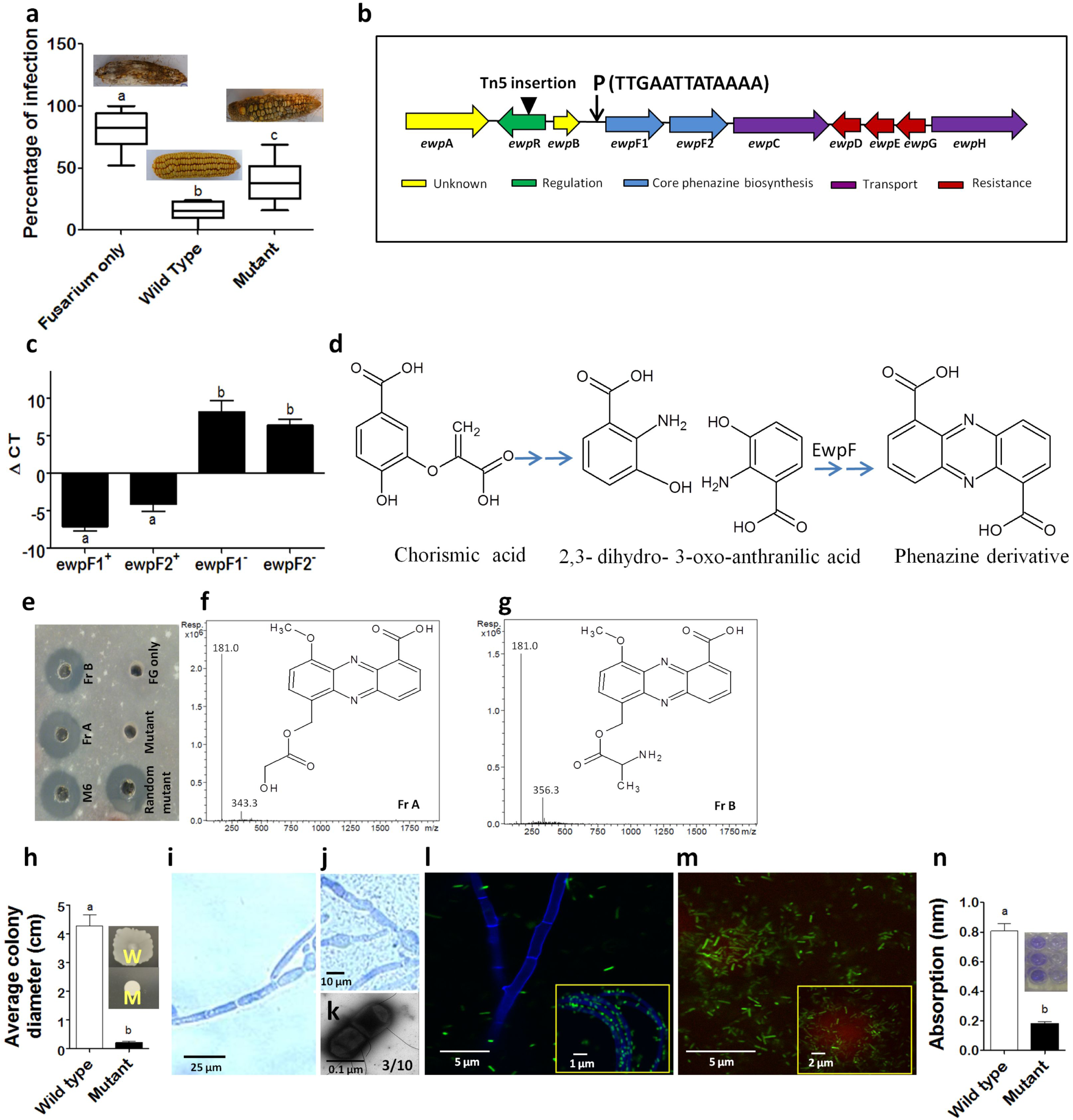
Characterization of phenazine mutant *ewpR-*5D7::Tn5. **a**, Effect of M6 mutant strain *ewpR-* on suppression of *F. graminearum* (Fg) in maize compared to wild type M6 and Fg-only control, with corresponding, representative maize ear pictures. **b**, Genomic organization of the predicted phenazine biosynthetic operon showing the position of the Tn5 insertion and putative LysR binding site within the promoter (P). **c**, Quantitative real time PCR (qRT-PCR) gene expression of the two core phenazine genes (*ewpF1* and *ewpF2*) in wild type M6 (+) and the mutant (-) (*ewpR-)*. **d**, Illustration of the phenazine biosynthetic pathway. **e**, Agar diffusion assay showing the inhibitory effect of different methanol extracts on the growth of Fg from wild type M6, two wild type fractions (FrA, FrB), the *ewpR-* mutant (mutant), a random Tn5 insertion or buffer. **f-g**, Mass spectroscopy analysis of putative phenazine derivatives in wild type M6 fractions A and B. **h**, Quantification of *ewpR-* mutant strain (M) motility compared to wild type M6 (W), with representative pictures (inset) of motility assays on semisolid agar plates. **i-j**, Light microscopy image showing loss of swarming and colony formation of (**i**) *ewpR-* mutant strain around Fg hyphae stained with lactophenol blue, compared to (**j**) wild type M6. **k**, Electron microscopy image of *ewpR-* mutant strain. **l**, Confocal microscopy image showing attachment pattern of GFP-tagged *ewpR-* mutant strain (green) to Fg hyphae stained with calcofluor stain, compared to wild type M6 (inset). **m**, Confocal microscopy image showing reduced proteinaceous biofilm matrix stained with Ruby film tracer (red) associated with GFP-tagged *ewpR-* mutant strain compared to wild type M6 (inset). **n**, Spectrophotometric quantification of biofilm formation associated with wild type M6 compared to the *ewpR-* mutant strain, with representative biofilm assay well pictures (left and right, respectively; 3 replicates shown). For graphs shown in (**a, c, h, n**) letters that are different from one another indicate that their means are statistically different (P≤0.05), and the whiskers represent the standard error of the mean.

Since M6, in the absence of *Fusarium*, was sporadically localized *in planta*, but then accumulated at sites of *Fusarium* hyphae, it was hypothesized that M6 actively seeks *Fusarium*. To test this hypothesis, GFP-tagged M6 and *F. graminearum* were spotted adjacent to one another on a microscope slide coated with agar; as time progressed, M6 was observed to swarm towards *Fusarium* hyphae (Fig. 3d-i), confirming its ability to seek *Fusarium.* Upon finding the pathogen, M6 cells were observed to physically attach onto *F. graminearum* hyphae (Fig. 3g-i). At the endpoint of these interactions, dense microcolonies of M6 were observed to break the hyphae (Fig. 3j). Transmission electron microscopy showed that M6 possesses multiple peritrichous flagella (Fig. 3k). Since this interaction was observed *in vitro*, independent of the host plant, the data show that M6 alone is sufficient to exert its fungicidal activity. To test if the attachments of M6 observed *in vitro* and *in planta* are mediated by biofilm formation, the proteinaceous biofilm matrix stain, Ruby Film Tracer (red), was used *in vitro*. Red staining, indicating biofilm formation, was observed associated with M6 in the absence of *Fusarium* (Fig. 3l). In the presence of *F. graminearum*, biofilm was also observed on the hyphal surfaces (Fig. 3m). Combined, these results suggest that M6 cells swarm towards *Fusarium* hyphae attempting to penetrate the root epidermis, induces root hair growth and bending, resulting in formation of RHESt within which M6 cells form biofilm-mediated microcolonies which attach, engulf and kill *Fusarium*.

### Identification of strain M6 genes required for anti-*Fusarium* activity

Since the fungicidal activity of M6 was observed to occur independently of its host plant, M6 was subjected to Tn5 mutagenesis and then candidate Tn5 insertions were screened *in vitro* for loss of fungicidal activity against *F. graminearum*. Out of 4800 Tn5 insertions that were screened in triplicate, sixteen mutants were isolated that resulted in loss or reduction in the diameter of inhibition zones of *F. graminearum* growth (Supplementary Fig. 5a). The mutants that resulted in complete loss of the antifungal activity *in vitro* were validated for loss of anti-*Fusarium* activity *in planta* in two independent greenhouse trials in maize (Supplementary Fig. 5b,c), demonstrating the relevance of the *in vitro* results. Rescue of the Tn5-flanking sequences followed by BLAST searching against the whole genome wild type M6 ^26^, resulted in the successful identification of 13 candidate genes in 12 predicted operons (Supplementary Table 5 and 6). Based on gene annotations and the published literature, four regulatory and/or anti-microbial mutants of interest were selected, complemented (Supplementary Fig. 5d) and subjected to detailed characterization. The selected genes encode two LysR family transcriptional regulators, a diguanylate cyclase, and a colicin V biosynthetic enzyme:

**Figure 5.**
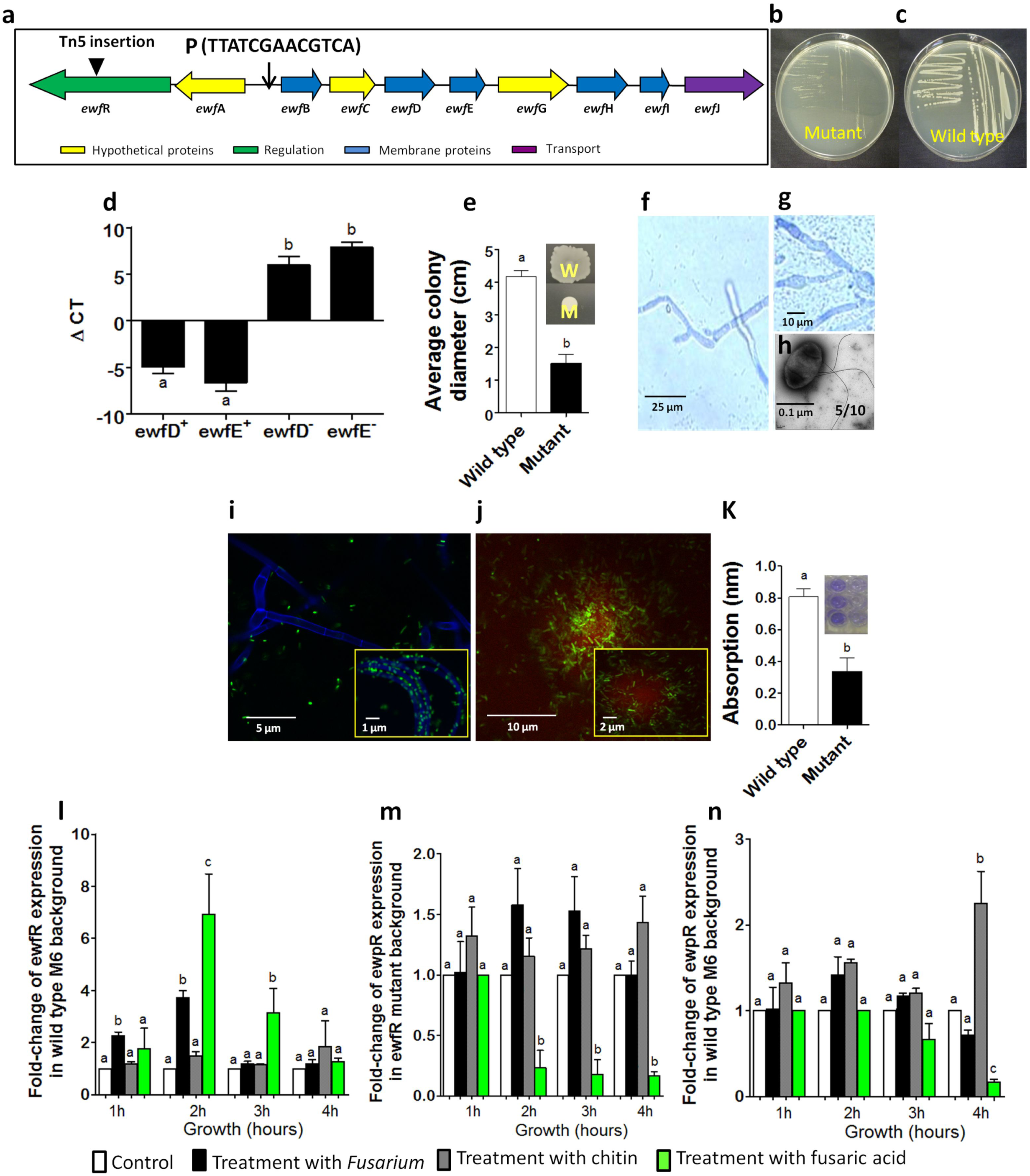
Characterization of fusaric acid resistance mutant *ewfR*-7D5::Tn5. **a**, Genomic organization of the predicted fusaric acid resistance operon showing the position of the Tn5 insertion and putative LysR binding site within the promoter (P). **b-c**, The inhibitory effect of fusaric acid embedded within agar on the growth of the *ewfR*- mutant compared to wild type M6. **d**, Quantitative real time PCR gene expression of two protein-coding genes required for the formation of the fusaric acid efflux pump (*ewfD* and *ewfE*) in wild type M6 (+) and the mutant (-) (*ewfR-*). **e**, Quantification of *ewfR-* mutant strain (M) motility compared to wild type M6 (W), with representative pictures (inset) of motility assays on semisolid agar plates. **f-g**, Light microscopy image showing decreased swarming and colony formation of (**f**) the *ewfR-* mutant strain around Fg hyphae stained with lactophenol blue, compared to (**g**) wild type M6. **h**, Electron microscopy image of the *ewfR-* mutant strain. **i**, Confocal microscopy image showing the attachment pattern of the GFP-tagged *ewfR-* mutant strain (green) to Fg hyphae stained with calcofluor stain, compared to wild type M6 (inset). **j**, Confocal microscopy image showing a proteinaceous biofilm matrix stained with Ruby film tracer (red) associated with GFP-tagged *ewfR-* mutant strain compared to wild type M6 (inset). **k**, Spectrophotometric quantification of biofilm formation associated with wild type M6 compared to the *ewfR-* mutant strain, with representative biofilm assay well pictures (left and right, respectively; 3 replicates shown). qRT PCR analysis of: (**l**) wild type *ewfR* expression in a wild type M6 background, (**m**) wild type *ewpR* in an *ewfR-* mutant background, and (**n**) wild type *ewpR* in a wild type M6 background. For graphs shown in (**d, e, k, l- n**) letters that are different from one another indicate that their means are statistically different (P≤0.05; in the case of **l-n**, within a time point), and the whiskers represent the standard error of the mean.

### LysR transcription regulator in a phenazine operon *(ewpR-*5D7::Tn5)

*Ewp-*5D7::Tn5 resulted in complete loss of the antifungal activity *in vitro* (Supplementary Fig. 5a) and reduction in activity *in planta* (Fig. 4a). The Tn5 insertion was localized to an operon (*ewp*, Fig. 4b) that included tandem paralogs of *phz*F (*ewp*F1 and *ewp*F1) (trans-2,3- dihydro-3-hydroxyanthranilate isomerase, EC # 5.3.3.17), a homodimer enzyme that forms the core skeleton of phenazine, a potent anti-fungal compound ^29^. The insertion occurred within a member of the LysR transcriptional regulator family (*ewpR*), which has been previously reported to induce phenazine biosynthesis ^30^. Three lines of evidence suggest the LysR gene is an upstream regulator of the *ewp* operon. First, the genomic organization showed that LysR was transcribed in the opposite direction as the operon. Second, the LysR canonical binding site sequence (TN11A) was observed upstream of the *phzF (ewaF)* coding sequences 31 (Fig. 4b). Finally, real time PCR revealed that the expression of *ewpF1* and *ewpF2* were dramatically down-regulated in the LysR mutant compared to wild type (Fig. 4c). Combined, these results suggest that EwpR is a regulator of phenazine biosynthesis in strain M6 (Fig. 4d). The crude methanolic extract from EwpR-5D7::Tn5 lost anti-*Fusarium* activity *in vitro* in contrast to extracts from wild type M6 or randomly selected Tn5 insertions that otherwise had normal growth rates (Fig. 4e). Anti-*F. graminearum* bioassay guided assay fractionation using extracts from M6 showed two active fractions (A, B) (Fig. 4e-g), each containing a compound with a diagnostic fragmentation pattern of [M+H]+=181.0, corresponding to a phenazine nucleus (C_12_H_8_N_2_, MW=180.08) ^32^, and molecular weights (M+Z= 343.3 and 356.3) indicative of phenazine derivatives [griseolutein A (C_17_H_14_N_2_O_6_, MW= 342.3) and D-alanyl-griseolutein (C_18_H_17_N_3_O_5_, MW=355.3), respectively] ^33-35^. Surprisingly, the *ewp* mutant showed a significant reduction in motility (Fig. 4h) and swarming (Fig. 4i) compared to the wild type (Fig. 4j), concomitant with a ∼60 % reduction in flagella (Fig. 4k) compared to wild-type (Fig. 3k), loss of attachment to *Fusarium* hyphae (Fig. 4l) compared to wild-type (inset in Fig. 4l), as well as reductions in biofilm formation (Fig. 4m,n). Combined, these results suggest that ewpR is required for multiple steps in the anti-fungal pathway of M6 including phenazine biosynthesis.

### LysR transcriptional regulator in a fusaric acid resistance pump operon (*ewf*R-7D5::Tn5)

*EwfR*-5D7::Tn5 resulted in a significant loss of the antifungal activity *in vitro* (Supplementary Fig. 5a). The Tn5 insertion occurred in an operon (*ewf*, Fig. 5a) that included genes that encode membrane proteins required for biosynthesis of the fusaric acid efflux pump including a predicted *fus*E-MFP/HIYD membrane fusion protein and *fus*E (*ewf*D and *ewf*E, respectively) and other membrane proteins (*ewfB*, *ewf*H and *ewf*I) ^36,37^. Fusaric acid (5-butylpyridine-2-carboxylic acid) is a mycotoxin that is produced by *Fusarium* which interferes with bacterial growth and metabolism and alters plant physiology ^38,39^. Bacterial-encoded fusaric acid efflux pumps promote resistance to fusaric acid ^40,41^. Consistent with expectations, *EwfR-*5D7::Tn5 failed to grow on agar supplemented with fusaric acid compared to the wild type (Fig. 5b,c). The Tn5 insertion specifically occurred within a member of the LysR transcriptional regulator family (*ewfR*). Similar to *ewpR* above and a previously published fusaric acid resistance operon ^41^, the regulator was transcribed in the opposite direction as the *ewf* operon, with this genomic organization suggesting that *ewfR* may be an upstream regulator of the operon. Indeed, the LysR canonical binding site sequence (TN_11_A) was observed upstream of the *ewfB-J* coding sequences ^31^ (Fig. 5a). Finally, real time PCR revealed that the expression of *ewfD* and *ewfE* were dramatically downregulated in the LysR mutant compared to wild type (Fig. 5d). Combined, these results suggest that EwfR is a positive regulator of the *ewf* fusaric acid resistance operon in strain M6.

In addition, the *ewf* mutant showed a significant reduction in motility (Fig. 5e) and swarming (Fig. 5f) compared to the wild type (Fig. 5g), concurrent with a ∼30 % reduction in flagella (Fig. 5h) compared to the wild-type (Fig. 3k). The mutant also showed loss of attachment to *Fusarium* hyphae (Fig. 5i) compared to wild-type (inset) and reductions in biofilm formation (Fig. 4j,k). Combined, these results suggest that strain M6 expression of resistance to fusaric acid is a pre-requisite step that enables subsequent anti-fungal steps.

### Interaction between the phenazine biosynthetic operon *(ewp)* and the fusaric acid resistance operon *(ewf)*

The expression of *ewf*R (LysR regulator of fusaric acid resistance) increased two-fold in the presence of *Fusarium* mycelium *in vitro* after 1 h of co-incubation, and tripled after 2 h (Fig. 5l). Expression of *ewfR* was also up-regulated by fusaric acid alone (Fig. 5l), demonstrating that the resistance operon is inducible. Fusaric acid has been shown to suppress phenazine biosynthesis through suppression of quorum sensing regulatory genes ^42^. Interestingly, expression of the putative LysR regulator of phenazine biosynthesis (*ewpR*, see above) was downregulated by fusaric acid at log phase (2.5-3 h), but only when fusaric acid resistance was apparently lost (*ewfR* mutant) (Fig. 5m) compared to wild type (Fig. 5n), suggesting that fusaric acid normally represses phenazine biosynthesis in M6. These results provide evidence for an epistatic relationship between the two LysR mutants required for the anti-*Fusarium* activity.

### Diguanylate cyclase (*ewg*S-10A8::Tn5)

*EwgS*-10A8::Tn5 resulted in a significant loss of the antifungal activity *in vitro* (Supplementary Fig. 5a). The Tn5 insertion occurred in a coding sequence encoding diguanylate cyclase (EC 2.7.7.65) (*ewgS*) that catalyzes conversion of 2-guanosine triphosphate to c-di-GMP, a secondary messenger that mediates quorum sensing and virulence traits ^43^ (Fig. 6a). Addition of exogenous c-di-GMP to the growth medium restored the antifungal activity of the mutant (Fig. 6b). Real time PCR showed that *ewgS* is not inducible by *Fusarium* (Fig. 6c). Consistent with its predicted upstream role in regulating virulence traits, the *ewg* mutant showed dramatic losses in motility (Fig. 6d), swarming (Fig. 6e, compared to wild-type Fig. 6f) and flagella formation (∼40 % reduction, Fig. 6g compared to wild-type, Fig. 3k). Attachment to *Fusarium* hyphae (Fig. 6h) and biofilm formation (Fig. 6i-j) appeared to have been lost completely.

**Figure 6.**
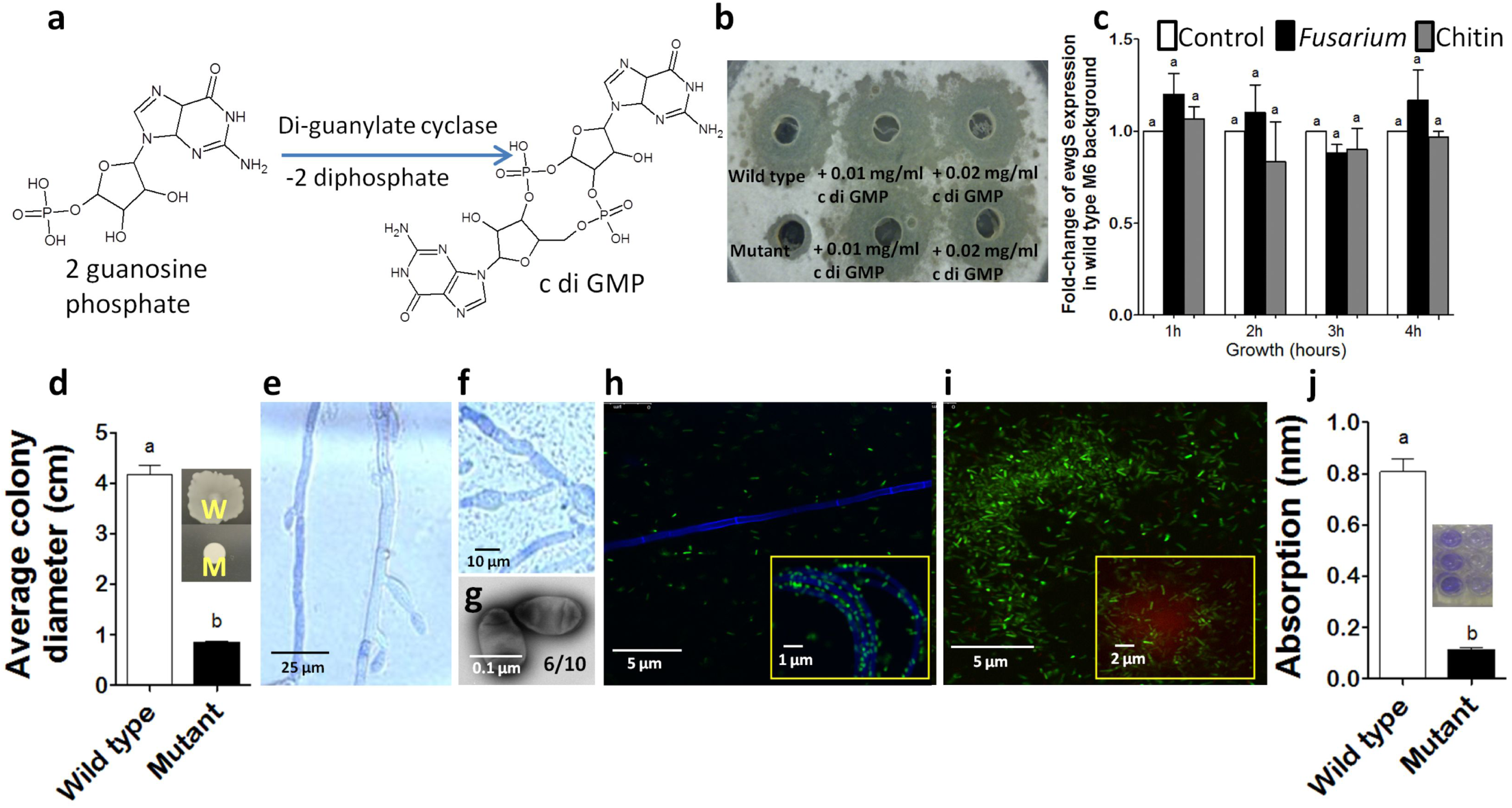
Characterization of di-guanylate cyclase mutant *ewgS*-10A8::Tn5. **a**, Illustration of the enzymatic conversion of 2 guanosine phosphate to c-di-GMP catalyzed by di-guanylate cyclase. **b**, Complementation of the putative *ewgS*- mutant with respect to inhibition of *F. graminearum* (Fg) by addition of c-di-GMP (0.01 and 0.02 mg/ml), compared to wild type strain M6. **c**, qRT-PCR analysis of wild type *ewgS* expression in a wild type M6 background. **d**, Quantification of *ewgS-* mutant strain (M) motility compared to wild type M6 (W), with representative pictures (inset) of motility assays on semisolid agar plates. **e-f**, Light microscopy image showing decrease in swarming and colony formation of (**e**) *ewgS-* mutant strain around Fg hyphae stained with lactophenol blue, compared to (**f**) wild type M6. **g**, Electron microscopy image of *ewgS-* mutant strain. **h**, Confocal microscopy image showing attachment pattern of GFP-tagged *ewgS-* mutant strain (green) to Fg hyphae stained with calcofluor stain, compared to wild type M6 (inset). **i**, Confocal microscopy image showing loss of proteinaceous biofilm matrix stained with Ruby film tracer (red) associated with GFP-tagged *ewgS-* mutant strain compared to wild type M6 (inset). **j**, Spectrophotometric quantification of biofilm formation associated with wild type M6 compared to the *ewgS-* mutant strain, with representative biofilm assay well pictures (left and right, respectively; 3 replicates shown). For graphs shown in (**c, d, j**) letters that are different from one another indicate that their means are statistically different (P≤0.05), and the whiskers represent the standard error of the mean.

### Colicin V production protein (*EwvC*-4B9::Tn5)

*EwvC*-4B9::Tn5 showed significant loss of the antifungal activity *in vitro* (Supplementary Fig. 5a). The Tn5 insertion occurred in a minimally characterized gene required for colicin V production (*ewvC*) orthologous to *cvpA* in *E.coli* ^44^. Colicin V is a secreted peptide antibiotic ^45^. Consistent with the gene annotation, *EwvC*-4B9::Tn5 failed to inhibit the growth of an *E. coli* strain that is sensitive to colicin V, compared to the wild type (Fig. 7a). Real time PCR showed that the expression of *ewvC* increased 6-fold after co-incubation with *Fusarium* at log phase (Fig. 7b). The mutant showed only limited changes in other virulence traits, including reductions in motility (Fig. 7c) and swarming (Fig. 7d, compared to wild-type, Fig. 7e), but with only a ∼15% minor reduction in flagella formation (Fig. 7f, compared to wild-type, Fig. 3k), no obvious change in attachment to *Fusarium* hyphae (Fig. 7g), and only a slight reduction in biofilm formation (Fig. 7h,i).

**Figure 7.**
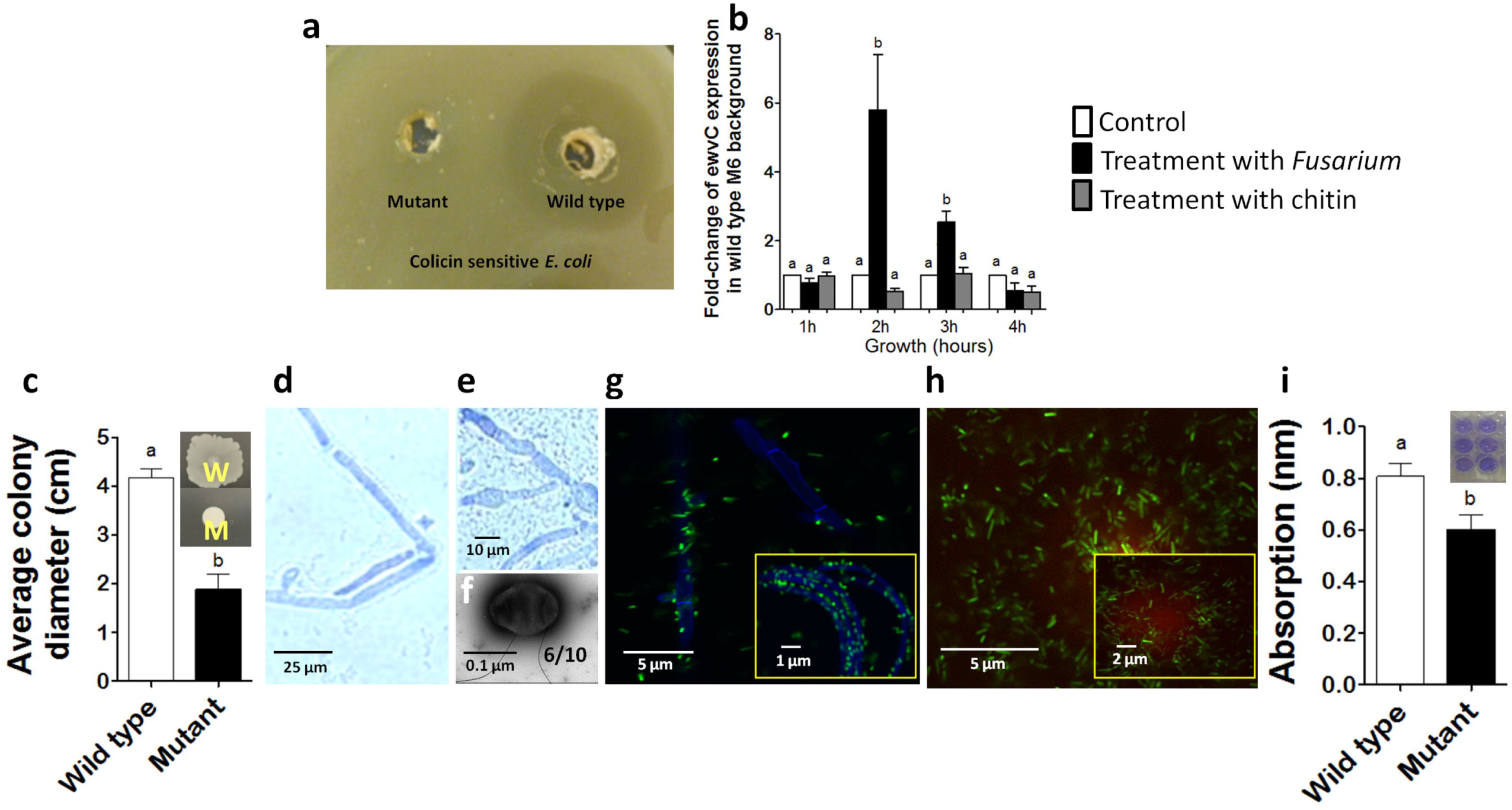
Characterization of Colicin V Mutant *ewvC*-4B9::Tn5. **a**, Dual culture agar diffusion assay showing loss of antagonism against the colicin V sensitive *E coli* strain (MC4100) by the *ewvC-* mutant compared to wild type M6. **b**, qRT-PCR analysis of wild type *ewvC* expression in a wild type M6 background. **c**, Quantification of *ewvC-* mutant strain (M) motility compared to wild type M6 (W), with representative pictures (inset) of motility assays on semisolid agar plates. **d-e**, Light microscopy image showing decrease in swarming and colony formation of (**d**) the *ewvC-* mutant strain around Fg hyphae stained with lactophenol blue, compared to (**e**) wild type M6. **f**, Electron microscopy image of the *ewvC-* mutant strain. **g**, Confocal microscopy image showing the attachment pattern of the GFP-tagged *ewvC-* mutant strain (green) to Fg hyphae stained with calcofluor stain, compared to wild type M6 (inset). **h**, Confocal microscopy image showing the proteinaceous biofilm matrix stained with Ruby film tracer (red) associated with the GFP-tagged *ewvC-* mutant strain compared to wild type M6 (inset). **i**, Spectrophotometric quantification of biofilm formation associated with wild type M6 compared to the *ewvC-* mutant strain, with representative biofilm assay well pictures (left and right, respectively; 3 replicates shown). For graphs shown in (**b, c, i**) letters that are different from one another indicate that their means are statistically different (P≤0.05), and the whiskers represent the standard error of the mean.

## Discussion

### RHESt as a novel plant defence mechanism

We hypothesized that the ancient African cereal finger millet hosts endophytic bacteria that contribute to its resistance to *Fusarium*, a pathogenic fungal genus that has been reported to share the same African origin as its plant target. Here, we report that a microbial inhabitant of finger millet (M6) actively swarms towards invading *Fusarium* hyphae, analogous to mobile immunity cells in animals, to protect plant cells that are immobile, confirming a hypothesis that we recently proposed ^46^. Endophyte M6 then builds a remarkable physico-chemical barrier resulting from root hair-endophyte stacking (RHESt) at the rhizosphere-root interface that prevents entry and/or traps *Fusarium* for subsequent killing. Mutant and biochemical data demonstrate that the killing activity of M6 requires genes encoding diverse regulatory factors, natural products and xenobiotic resistance. The RHESt consists of two lines of defence, a dense layer of intercalated root hairs and endophyte microcolonies followed by a long, continuous endophyte barrier layer on the root epidermal surface (see summary model, Supplementary Fig. 6). RHESt represents a plant defence mechanism that has not been previously captured to the best of our knowledge and is an unusual example of host-microbe symbiosis.

The epidermal root surface where microbes reside is termed the rhizoplane ^47-51^. Soil microbes have previously been reported to form biofilm-mediated aggregates on the rhizoplane ^52-57^ sometimes as part of their migration from the soil to the root endosphere ^48,58^ and may prevent pathogen entry ^48,59^. However, here we report that an endophyte, not soil microbe, forms a pathogen barrier on the rhizoplane. The previously reported soil microbes on the rhizoplane are thought to take advantage of nutrient-rich root exudates ^59^, whereas RHESt is a *de novo*, inducible structure that only forms in the presence of *Fusarium* in coordination with root hairs.

M6 creates its own specialized killing microhabitat by inducing growth of local root hairs which are then bent to form the RHESt scaffold, likely mediated by biofilm formation and attachment. *In vitro*, we observed that M6 synthesizes auxin (IAA), a hormone known to stimulate root hair growth ^60^ and that can be synthesized by microbes ^61^. Root hair bending associated with RHESt might be an active process, similar to rhizobia-mediated root hair curling ^62^, or an indirect consequence of micro-colonies attachment to adjacent root hairs.

### Regulatory signals within the anti-fungal pathway

Some of the mutants caused pleiotropic phenotypes, including loss of swarming, attachment and/or biofilm formation, which was a surprising result. These mutants included the transcription factors associated with operons for phenazine biosynthesis, fusaric acid resistance, as well as formation of c-di-GMP. One interpretation of these results is that the underlying genes help to regulate the early steps of the anti-fungal RHESt pathway. Indeed, phenazines have been reported to act as signalling molecules that regulate the expression of hundreds of genes including those responsible for motility and defense ^63^. Our results showed an epistatic relationship between the two transcription factors regulating phenazine biosynthesis and fusaric acid resistance, with the latter required for the former (Fig. 5m,n), suggesting that the pleiotropic phenotypes observed in the fusaric acid resistance regulatory mutant may have been mediated by a reduction in phenazine signaling. Finally, c-di-GMP as a sensor of the environment and population density ^64,65^, and a secondary messenger ^66^ involved in transcriptional regulation of genes encode virulence traits such as motility, attachment, and biofilm formation^43,64,66-68^, all activities that would be logically required for the RHESt-mediated anti-fungal pathway. Genome mining of strain M6 also revealed the presence of a biosynthetic cluster for 2, 3 butanediol which is a hormone known to elicit plant defences ^69^. Production of 2, 3 butanediol was confirmed by LC-MS analysis (Supplementary Fig. 7a). Butanediol is thus a candidate signalling molecule for M6-millet cross-talk.

### Fungicidal compounds required for M6 killing activity

In addition to RHESt formation, we gained evidence that bacterial endophyte M6 evolved multiple biochemical strategies to actively break and kill *Fusarium* hyphae (Fig. 2j and Fig. 3a,j) involving diverse classes of natural products, including phenazines, colicin V, chitinase and potentially other metabolites.

*Phenazines:* Phenazines are heterogeneous nitrogenous compounds produced exclusively by bacteria ^70^. Phenazines exhibit potent antifungal activity, in particular against soil pathogens including *Fusarium* species ^71-73^. Here M6 was observed to produce at least two distinct phenazines, D-alanyl-griseolutein and griseolutein A which previously shown to have anti-microbial activities ^74^. Phenazines lead to the accumulation of reactive oxygen species in target cells, due to their redox potential ^75,76^. Phenazines were also reported to induce host resistance ^77^. For bacteria to survive inside a biofilm, where oxygen diffusion is limited, molecules with high redox potential such as phenazines are required ^78-82^. Furthermore, the redox reaction of phenazine releases extracellular DNA which enhances surface adhesion and cellular aggregation of bacteria to form a biofilm ^83-88^. Consistent with these roles, phenazines are known to be produced inside biofilms ^78^, and are required for biofilm formation ^89^, which might explain the diminished biofilm formation observed with the phenazine-associated LysR mutant. Since M6 was observed to produce biofilm around its fungal target (Fig. 3m) which is killed within RHESt (Fig. 2j), phenazines may be part of the killing machinery.

*Colicin V:* An exciting observation from this study is that colicin V also appears to be required for the fungicidal activity of strain M6 (Fig. 7). Colicin V is a small peptide antibiotic belonging to the bacteriocin family which disrupts the cell membrane of pathogens resulting in loss of membrane potential ^90^. Bacteriocins are generally known for their antibacterial activities ^91^. Indeed, we could find only one previous report of colicin V having anti-fungal activity ^92^. Our results suggest that this compound may target a wider spectrum of pathogens than previously thought.

*Chitinase*: A Tn5 insertion in a gene (*ewc-*3H2::Tn5)encodes chitinase (EC 3.2.1.14), also resulted in a significant loss of the antifungal activity (Supplementary Fig. 5a). The mutant shows a significant reduction in production of chitinase *in vitro*, compared to wild type M6 (Supplementary Fig. 7b). Chitinase exerts its antifungal activity by hydrolyzing chitin, a principle component of the fungal cell wall ^93,94^.

*Other putative M6 anti-fungal metabolites:* Two additional putative Tn5 mutants suggest that other metabolites may be involved in the anti-*Fusarium* activity of strain M6 within the RHESt, specifically phenylacetic acid (PAA) and P-amino-phenyl-alanine antibiotics (PAPA). The requirement for PAA was suggested by a mutant in phenylacetic acid monoxygenase which resulted in loss of anti-*Fusarium* activity (m2D7, Supplementary Fig. 5a). This enzyme catalyzes the biosynthesis of hydroxy phenylacetic acid, derivatives of which have been shown to act as anti-fungal compounds ^95,96^. In an earlier report, an *Enterobacter* sp. that was used to control Fusarium dry rot during seed storage ^97^ was shown to require phenylacetic acid, indole-3-acetic acid (IAA) and tyrosol ^96^. In addition to PAA implicated here by the Tn5 mutant, wild type strain M6 was shown to produce IAA *in vitro* (Supplementary Fig. 4).

The requirement for PAPA was suggested by a putative mutant (m15A12, Supplementary Fig. 5a) which disrupted a gene encoding a permease transport protein that is a part of an operon responsible for biosynthesis of PAPA and 3-hydroxy anthranilates. PAPA is the direct precursor of well known antibiotics including chloramphenicol and obafluorin ^98-100^.

### M6-*Fusarium*-millet co-evolution and the fusaric acid-phenazine arms race

Molecular and biochemical data suggest that the anti-fungal activity of M6 requires diverse classes of anti-fungal natural products (phenazine metabolites, colicin V peptide antibiotic, chitinase enzyme). We previously demonstrated that finger millet also hosts fungal endophytes that secrete complementary anti-*F. graminearum* natural products including polyketides and alkaloids ^4^. These observations, combined with loss of function mutants from this study that demonstrate that no single anti-fungal mechanism is sufficient for M6 to combat *Fusarium*, suggests that the endophytic community of finger millet and *Fusarium* have been engaged in a step-by-step arms race that resulted in the endophytes having a diverse weapons arsenal, presumably acting within RHESt. Consistent with this interpretation, mutant analysis showed that the anti-fungal activity of M6 requires a functional operon that encodes resistance to the *Fusarium* mycotoxin, fusaric acid. Furthermore, our results show a novel epistatic regulatory interaction between the fusaric acid resistance and phenazine pathways, wherein an M6-encoded LysR activator of fusaric acid resistance prevents fusaric acid from suppressing expression of the M6-encoded LysR regulator of phenazine biosynthesis (Fig. 5m). Fusaric acid has previously been shown to interfere with quorem sensing-mediated biosynthesis of phenazine ^42^. We propose that the phenazine-fusaric acid arms race provides a molecular and biochemical paleontological record that M6 and *Fusarium* co-evolved.

We show how this tripartite co-evolution likely benefits subsistence farmers not only by suppressing *Fusarium* entry and hence disease in plants, but also in seeds after harvest. Specifically, under poor seed storage conditions that mimic those of subsistence farmers, M6 caused dramatic reductions in contamination with DON (Fig. 1m,r), a potent human and livestock mycotoxin. Hence, in the thousands of years since ancient crop was domesticated, farmers may have been inadvertently selecting for the physico-chemical RHESt barrier activity of endophyte M6, simply by selecting healthy plants and their seeds. We have shown here that the benefits of M6 are transferable to two of the world’s most important modern crops, maize and wheat (Fig. 1i,o), which are severely afflicted by *F. graminearum* and DON. In addition to *F. graminearum*, M6 inhibited the growth of five other fungi including two additional *Fusarium* species (Supplementary Table 2), suggesting that RHESt-mediated plant defence may contribute to the broad spectrum pathogen resistance of finger millet reported by subsistence farmers. Despite its importance, finger millet is a scientifically neglected crop ^1,2^. Our study suggests the value of exploring microbiome--host interactions of other scientifically neglected, ancient crops.

## Methods

### Isolation, identification and antifungal activity of endophytes

Finger millet seeds originating from Northern India were grown on clay Turface in the summer of 2012 according to a previously described method ^4^. Tissue pool sets (3 sets of: 5 seeds, 5 shoots and 5 root systems from pre-flowering plants) were surface sterilized following a standard protocol ^4^. Surface sterilized tissues were ground in LB liquid medium in a sterilized mortar and pestle, then 50 µl suspensions were plated onto 3 types of agar plates (LB, Biolog Universal Growth, and PDA media). Plates were incubated at 25°C, 30°C and 37°C for 1-3 days. A total of seven bacterial colonies (M1-M7) were purified by repeated culturing on fresh media. For molecular identification of the isolated bacterial endophytes, PCR primers (Supplemental Table 7) were used to amplify and sequence 16S rDNA as previously described ^7^, followed by best BLAST matching to GenBank. 16S rDNA sequences were deposited into GenBank. Scanning electron microscopy imaging was used to phenotype the candidate bacterium as previously described ^7^ using a Hitachi S-570 microscope (Hitachi High Technologies, Tokyo, Japan) at the Imaging Facility, Department of Food Science, University of Guelph.

To test the antifungal activity of the isolated endophytes against *F. graminearum*, agar diffusion dual culture assays were undertaken in triplicate ^101^. Nystatin (Catalog #N6261, Sigma Aldrich, USA) and Amphotericin B (Catalog #A2942, Sigma Aldrich, USA) were used as positive controls at 10 µg/ml and 5 µg/ml, respectively, while the negative control was LB medium. Using a similar methodology, additional anti-fungal screening was conducted using the Fungal Type Culture Collection at Agriculture and Agrifood Canada, Guelph, ON (Supplementary Table 2).

### Microscopy imaging

#### *In planta* colonization of the candidate anti-fungal endophyte M6

In order to verify the endophytic behaviour of the candidate anti-fungal bacterium M6 in maize, wheat and millet, the bacterium was subjected to tagging with green fluorescent protein (GFP) (vector pDSK-GFPuv) ^101^ and *in planta* visualization using confocal scanning microscopy as previously described ^101^ at the Molecular and Cellular Imaging Facility, University of Guelph, Canada.

#### *In vitro* interaction using light microscopy

Both the fungus and bacterium M6 were allowed to grow in close proximity to each other overnight on microscope slides coated with a thin layer of PDA as previously described ^101^. Thereafter, the fungus was stained with the vitality stain, Evans blue which stains dead hyphae blue. The positive control was a commercial biological control agent (*Bacillus subtilis* QST*713,* Bayer CropScience, Batch # 00129001) (100 mg/10 ml). Pictures were captured using a light microscope (BX51, Olympus, Tokyo, Japan).

#### *In vitro* and *in planta* interactions using confocal microscopy

All the experiments were conducted using a Leica TCS SP5 confocal laser scanning microscope at the Molecular and Cellular Imaging Facility at the University of Guelph, Canada.

To visualize the interactions between endophyte M6 and *F. graminearum* inside finger millet, finger millet seeds were surface sterilized and coated with GFP-tagged endophyte, and then planted on sterile Phytagel based medium in glass tubes, each with 4-5 seeds. Phytagel medium was prepared as previously described ^4^. At 14 days after planting, finger millet seedlings were inoculated with *F. graminearum* (50 µl of a 48 h old culture grown in potato dextrose broth) and incubated at room temperature for 24 h. The control consisted of seedlings incubated with potato dextrose broth only. There were three replicate tubes for each treatment. Thereafter, roots were stained with calcofluor florescent stain (catalog #18909, Sigma-Aldrich), which stains chitin blue, following the manufacturer’s protocol, and scanned.

To visualize the attachment of bacterium M6 to fungal hyphae, GFP-tagged M6 and *F. graminearum* were inoculated overnight at 30°C on microscope slides covered with a thin layer of PDA. Thereafter, the fungal hyphae was stained with calcofluor and examined. The same protocol was employed to test if this recognition was disrupted in the Tn5 mutants.

To visualize biofilm formation by bacterium M6, GFP-tagged M6 were incubated on microscope slides for 24 h at 30°C. The biofilm matrix was stained with FilmTracer™ SYPRO® Ruby Biofilm Matrix Stain (F10318) using the manufacturer’s protocol and then examined. The same protocol was employed to test if the biofilm was disrupted in Tn5 mutants.

### Suppression of *F. graminearum in planta* and accumulation in storage

Bacterium M6 was tested *in planta* for its ability to suppress *F. graminearum* in two susceptible crops, maize (hybrid P35F40, Pioneer HiBred) and wheat (Quantum spring wheat, C&M Seeds, Canada), in two independent greenhouse trials as previously described for maize ^101^, with modifications for wheat (Supplemental Method 2). ELISA analysis was conducted to test the accumulation of DON in seeds after 14 months of storage in conditions that mimic those of African subsistence farmers (temperature ∼18-25°C, with a moisture content of ∼40-60%) as previously described ^101^. Control treatments consisted of plants subjected to pathogen inoculation only, and plants subjected to pathogen inoculation followed by prothioconazole fungicide spraying (PROLINE® 480 SC, Bayer Crop Science). Results were analyzed and compared using Mann-Whitney t-tests (P≤0.05).

### Transposon mutagenesis, gene rescues and complementation

To identify the genes responsible for the antifungal activity, Tn5 transposon mutagenesis was conducted using EZ-Tn5 <R6Kγori/KAN-2>Tnp Transposome^TM^ kit (catalog #TSM08KR, Epicentre, Madison, USA) according to the manufacturer’s protocol. The mutants were screened for loss of anti-*Fusarium* activity using the agar dual culture method compared to wild type. Insertion mutants that completely lost the antifungal activity *in vitro* were further tested for loss of *in planta* activity using maize as a model in two independent greenhouse trials (same protocol as described above). The sequences flanking each candidate Tn5 insertion mutant of interest were identified using plasmid rescues according to the manufacturer’s protocol (Supplemental Method 3). The rescued gene sequences were BLAST searched against the whole genome sequence of bacterium M6 ^26^. To test if the candidate genes are inducible by *F. graminearum* or constitutively expressed, real-time PCR analysis was conducted using gene specific primers (Supplemental Method 4). Operons were tentatively predicted using FGENESB ^102^ from Softberry Inc. (USA). Promoter regions were predicted using PePPER software (University of Groningen, The Netherlands) ^103^. In order to confirm the identity of the genes discovered by Tn5 mutagenesis, each mutant was complemented with the corresponding predicted wild type coding sequence which was synthesized (VectorBuilder, Cyagen Biosciences, USA) using a pPR322 vector backbone (Novagen) under the control of the T7 promoter. Two microlitres of each synthesized vector was electroporated using 40 µl electro competent cells of the corresponding mutant. The transformed bacterium cells were screened for gain of the antifungal activity against *F. graminearum* using the dual culture assay as described above.

### Mutant phenotyping

*Transmission electron microscopy (TEM):* To phenotype the candidate mutants, TEM imaging was conducted. Wild type strain M6 and each of the candidate mutants were cultured overnight in LB medium (37°C, 250 rpm). Thereafter, 5 µl of each culture were pipetted onto a 200-mesh copper grid coated with carbon. The excess fluid was removed onto a filter, and the grid was stained with 2% uranyl acetate for 10 sec. Images were taken by a F20 G2 FEI Tecnai microscope operating at 200 kV equipped with a Gatan 4K CCD camera and Digital Micrograph software at the Electron Microscopy Unit, University of Guelph, Canada.

*Motility assay:* Wild type or mutant strains were cultured overnight in LB medium (37°C, 225 500 rpm). The OD_595_ for each culture was adjusted to 1.0, then 15 µl of each culture were spotted on the center of a semisolid LB plates (0.3% agar) and incubated overnight (37°C, no shaking). Motility was measured as the diameter of the resulting colony. There were ten replicates for each culture. The entire experiment was repeated independently.

*Swarming test:* To examine the ability of the strains to swarm and form colonies around the fungal pathogen, *in vitro* interaction/light microscopy imaging was conducted. Wild type M6 and each mutant were incubated with *F. graminearum* on microscope slides covered with PDA (as described above). *F. graminearum* hyphae was stained with lactophenol blue solution (catalog #61335, Sigma-Aldrich) then examined under a light microscope (B1372, Axiophot, Zeiss, Germany) using Northern Eclipse software.

*Biofilm spectroscopic assay:* To test the ability of the strains to form biofilms, the strains were initially cultured overnight in LB medium (37°C and 250 rpm), and adjusted to OD_600_ of 0.5. Cultures were diluted in LB (1:100), thereafter, 200 µl from each culture were transferred to a 96 well microtitre plate (3370, Corning Life Sciences, USA) in 6 replicates. The negative control was LB medium only. The microtitre plate was incubated for 2 days at 37°C. The plate was emptied by aspiration and washed three times with sterile saline solution. Adherent cells were fixed with 200 µl of 99% methanol for 15 min then air dried. Thereafter, 200 µl of 2% crystal violet (94448, Sigma) were added to each well for 5 min then washed with water, and left to air dry. Finally, 160 µl of 33% (v/v) glacial acetic acid were added to each well to solubilise the crystal violet stain. The light absorption was read by a spectrophotometer (SpectraMax plus 348 microplate reader, Molecular Devices, USA) at 570 nm ^104^. The entire experiment was repeated independently.

*Fusaric acid resistance:* To test the ability of M6 wild type or *EwfR::Tn5* to resist fusaric acid, the strains were allowed to grow on LB agar medium supplemented with different concentrations of fusaric acid (0.01, 0.05 and 0.1%, catalog #F6513, Sigma Aldrich) as previously reported ^105^.

*c-di-GMP chemical complementation:* To test if the Tn5 insertion in the predicted guanylate cyclase gene could be complemented by exogenous c-di-GMP, M6 wild type or *EwgC::Tn5* strains were grown on LB agar medium supplemented with c-di-GMP (0.01 or 0.02 mg/ml) (Cataloge # TLRL-CDG, Cedarlane). After 24 h, the agar diffusion method was used to test for antifungal activity.

*Colicin V assay:* To verify that the Tn5 insertion in the predicted colicin V production gene caused loss of colicin V secretion, M6 wild type or *EwvC*-4B9::Tn5 strains were inoculated as liquid cultures (OD_600_=0.5) into holes created in LB agar medium pre-inoculated after cooling with an *E.coli* strain sensitive to colicin V (MC4100, ATCC 35695) ^90^.

### Biochemical and enzymatic assays

*Detection of anti-fungal phenazine derivatives:* For phenazine detection, bio-guided fractionation combined with LC-MS analysis was undertaken. Wild type M6 or mutant bacterial strains were grown for 48 h on Katznelson and Lochhead liquid medium ^106^, harvested by freeze drying, then the lyophilized powder from each strain was extracted by methanol. The methanolic extracts were tested for anti-*Fusarium* activity using the agar diffusion method as described above. The extracts were dried under vacuum and dissolved in a mixture of water and acetonitrile then fractionized over a preparative HPLC C18 column (Nova-Pak HR C18 Prep Column, 6 µm, 60 Å, 25 x 100 mm prepack cartridge, part # WAT038510, Serial No 0042143081sp, Waters Ltd, USA). The solvent system consisted of purified water (Nanopure, USA) and acetonitrile (starting at 99:1 and ending at 0:100) pumped at a rate of 5 ml/min. The eluted peaks were tested for anti-*Fusarium* activity. Active fractions were subjected to LC-MS analysis. Each of the active fractions was run on a Luna C18 column (Phenomenex Inc, USA) with a gradient of 0.1% formic acid in water and 0.1% formic in acetonitrile. Peaks were analyzed by mass spectroscopy (Agilent 6340 Ion Trap), ESI, positive ion mode. LC-MS analysis was conducted at the Mass Spectroscopy Facility, McMaster University, ON, Canada.

*GC-MS to detect production of 2,3 butanediol:* To detect 2,3 butanediol, wild type strain M6 was grown for 48 h on LB medium. The culture filtrate was analyzed by GC-MS (Mass Spectroscopy Facility, McMaster University, ON, Canada) and the resulting peaks were analyzed by searches against the NIST 2008 database.

*Chitinase assay:* Chitinase activity of wild type strain M6 and the putative chitinase Tn5 mutant was assessed using a standard spectrophotometric assay employing the Chitinase Assay Kit (catalog #CS0980, Sigma Aldrich, USA) according to the manufacturer’s protocol. There were three replicates for each culture, and the entire experiment was repeated independently.

## Supplementary Data

**Supplementary Table 1.** Taxonomic classification of finger millet bacterial endophytes based on 16S rDNA sequences and BLAST analysis.

| **ID (Genbank Accession)** | **Tissue source** | **BLAST analysis** | **% of max. identity** | **Length of aligned sequence** |
| --- | --- | --- | --- | --- |
| M1(KU307449) | Roots | *Enterobacter* sp. (CP015227) | 99 | 646 |
| M2 (KU307450) | Seeds | *Pantoea* sp. (FN796868) | 99 | 701 |
| M3(KU307451) | Seeds | *Burkholderia* sp. (KC522298) | 99 | 587 |
| M4 (KU307452) | Shoots | *Pantoea* sp. (KT075171) | 95 | 751 |
| M5 (KU307453) | Shoots | *Burkholderia* sp. (KP455296) | 99 | 586 |
| M6 KU307454 | Roots | *Enterobacter* sp. (KU935452) | 99 | 586 |
| M7 (KU307455) | Shoots | *Burkholderia* sp. (HQ023278) | 100 | 644 |

**Supplementary Table 2.** Effect of endophyte strain M6 isolated from finger millet on the growth of diverse fungal pathogens *in vitro*.

| Target fungal species | Diameter of growth inhibition zone in mm |  |  |
| --- | --- | --- | --- |
|  | Nystatin<br>(10.0 U/ml) | Amphotericin<br>(250 µg/ml) | M6 |
| *Alternaria alternata* | 0.0±0.0 | 0.0±0.0 | 0.0±0.0 |
| *Alternaria arborescens* | 0.0±0.0 | 0.0±0.0 | 0.0±0.0 |
| *Aspergillus flavus* | 2.0±0.2 | 0.0±0.0 | 3.6±0.2 |
| *Aspergillus niger* | 0.0±0.0 | 2.0±0.0 | 0.0±0.0 |
| *Bionectria ochroleuca* | 2.0±0.2 | 0.5±0.2 | 0.0±0.0 |
| *Davidiella tassiana* | 1.5±0.2 | 0.5±0.3 | 0.0±0.0 |
| *Diplodia pinea* | 2.5±0.2 | 3.0±0.2 | 0.0±0.0 |
| *Diplodia seriata* | 3.0±0.2 | 2.0±0.2 | 0.0±0.0 |
| *Epicoccum nigrum* | 0.0±0.0 | 0.0±0.0 | 0.0±0.0 |
| *Fusarium avenaceum* (isolate 1) | 2.5±0.3 | 3.0±0.6 | 1.8±0.2 |
| *Fusarium graminearum* | 1.5±1.6 | 0.0±0.0 | 5.0±0.3 |
| *Fusarium lateritium* | 0.0±0.0 | 1.0±0.2 | 0.0±0.0 |
| *Fusarium sporotrichioides* | 1.0±0.2 | 1.0±0.2 | 2.8±0.2 |
| *Fusarium avenaceum* (isolate 2) | 0.0±0.0 | 0.0±0.0 | 0.0±0.0 |
| *Nigrospora oryzae* | 0.0±0.0 | 0.0±0.0 | 1.83±0.2 |
| *Nigrospora sphaerica* | 0.0±0.0 | 0.0±0.0 | 0.0±0.0 |
| *Paraconiothyrium brasiliense* | 0.0±0.0 | 0.0±0.0 | 0.0±0.0 |
| *Penicillium afellutanum* | 3.0±0.2 | 3.0±0.2 | 0.0±0.0 |
| *Penicillium expansum* | 2.0±0.2 | 5.0±0.2 | 4.9±0.1 |
| *Penicillium olsonii* | 1.5±0.3 | 3.5±0.3 | 0.0±0.0 |
| *Rosellinia corticium* | 2.0±0.2 | 4.5±0.3 | 0.0±0.0 |

**Supplementary Table 3.** Suppression of *F. graminearum* disease symptoms in maize and wheat by endophyte M6 in replicated greenhouse trials.

| **Treatment** | **% Infection**<br>**(mean ± SEM)\*** | **% Disease reduction relative to *Fusarium* only treatment** | **Average yield per plant\*** | **% Yield increase relative to *Fusarium* only treatment** |
| --- | --- | --- | --- | --- |
| **Greenhouse trial 1 in maize** |  |  |  |  |
| *Fusarium* only | 33.69±2.3 | 0.0 | 41.77±1.4 | 0.0 |
| Proline fungicide | 23.10±2.0 | 31.4 | 50.9±1.2 | 21.85 |
| M6 | 10.13±0.9 | 69.93 | 39.58±2.6 | -5.2 |
| **Greenhouse trial 2 in maize** |  |  |  |  |
| *Fusarium* only | 85.11±4.5 | 0.0 | 7.8±1.1 | 0.0 |
| Proline fungicide | 41.42±4.5 | 51.33 | 40.8±1.2 | 423 |
| M6 | 6.9±0.9 | 91.89 | 34.70±2.7 | 344 |
| **Greenhouse trial 1 in wheat** |  |  |  |  |
| *Fusarium* only | 51.8±6.3 | 0.0 | 10.5±2.6 | 0.0 |
| Proline fungicide | 31.70±2.6 | 38.8 | 10.9±1.3 | 3.8 |
| M6 | 41.1±2.6 | 20.6 | 10.6±0.8 | 0.95 |
| **Greenhouse trial 2 in wheat** |  |  |  |  |
| *Fusarium* only | 54.0±2.6 | 0.0 | 17.7±1.5 | 0.0 |
| Proline fungicide | 31.5±4.1 | 41.6 | 20.1±2.1 | 13.5 |
| M6 | 30.7±3.6 | 43.1 | 21.1±1.3 | 19.2% |

**\*SEM is the standard error of the mean.**

**Supplementary Table 4.** Reduction of DON mycotoxin accumulation during prolonged seed storage following treatment with endophyte M6.

| **Treatment** | **DON content (ppm)**<br>**(mean ± SEM)\*** | **% DON reduction relative to**<br>**Fusarium only treatment\*** |
| --- | --- | --- |
| **Greenhouse trial 1 in maize** |  |  |
| *Fusarium* only | 3.4±0.4 | 0.0 |
| Proline fungicide | 0.7±0.4 | 79.4 |
| M6 | 0.1±0.0 | 97 |
| **Greenhouse trial 2 in maize** |  |  |
| *Fusarium* only | 3.5±0.3 | 0.0 |
| Proline fungicide | 0.1±0.0 | 97.1 |
| M6 | 0.1±0.0 | 97.1 |
| **Green house trial 1 in wheat** |  |  |
| *Fusarium* only | 7.6±0.3 | 0.0 |
| Proline fungicide | 0.5±0.1 | 94.6 |
| M6 | 1.5±0.6 | 81.33 |
| **Green house trial 2 in wheat** |  |  |
| *Fusarium* only | 5.5±0.7 | 0.0 |
| Proline fungicide | 1.3±0.9 | 76.3 |
| M6 | 2.0±0.4 | 63.6 |

**\*SEM is the standard error of the mean.**

**Supplementary Table 5.** Complete list of strain M6 Tn5 insertion mutants showing loss of antifungal activity against *F. graminearum in vitro*.

| ID | Gene prediction | Swarming assay | Motility assay (mean ± SEM)* | Presence of flagella (%) | Biofilm (mean ± SEM)* |
| --- | --- | --- | --- | --- | --- |
| Wild type | | +++ | 4.2±0.3 | 70% | 0.8 |
| *ewa-1B3::Tn5*<br>*ewa-4B8::Tn5* | Transcription regulator, AraC | --- | 0.6±0.02 | 50% | 0.11±0.00 |
| *ewy-1C5::Tn5*<br>*ewy-5D2::Tn5* | YjbH outer-membrane protein | ++ | 0.5±0.03 | 10% | 0.3±0.08 |
| *ewm-2D7::Tn5* | 4-hydroxyphenyl acetate 3-monooxygenase<br>May catalyze production of phenylacetic acid (PAA) | --- | 0.8±0.05 | 20% | 0.07±0.00 |
| *ewpR-5D7::Tn5* | Transcription regulator, LysR | --- | 0.5±0.03 | 30% | 0.18±0.01 |
| *ewb-9F12::Tn5*<br>*ewb-7C5::Tn5* | Fatty acid biosynthesis | --- | 0.6±0.05 | 30% | 0.3±0.11 |
| *ewvC-4B9::Tn5* | Colicin V production | + | 1.8±0.30 | 60% | 0.6±0.05 |
| *ewT-15A12::Tn5* | Transport permease protein<br>Within operon for biosynthesis of P-amino-phenyl-alanine antibiotics (PAPA). | + | 1.5±0.00 | 30% | 0.6±0.04 |
| *ews-1H3::Tn5* | Sensor histidine kinase | + | 0.8±0.16 | 50% | 0.8±0.2 |
| *ewc-3H2::Tn5* | Chitinase | --- | 1.0±0.00 | 20% | 0.4±0.15 |
| *ewfR-7D5::Tn5* | Transcription regulator, LysR | + | 1.5±0.20 | 50% | 0.3±0.08 |
| *ewgS-10A8::Tn5* | Di-guanylate cyclase | --- | 0.8±0.30 | 40% | 0.1±0.01 |
| *ewh-5B1::Tn5* | Hig A protein | +++ | 1.6±0.30 | 30% | 0.6±0.10 |
| *ewa-8E1::Tn5* | Hypothetical protein | ++ | 2.3±0.40 | 20% | 0.5±0.09 |

\*SEM is the standard error of the mean.

**Supplementary Table 6.**
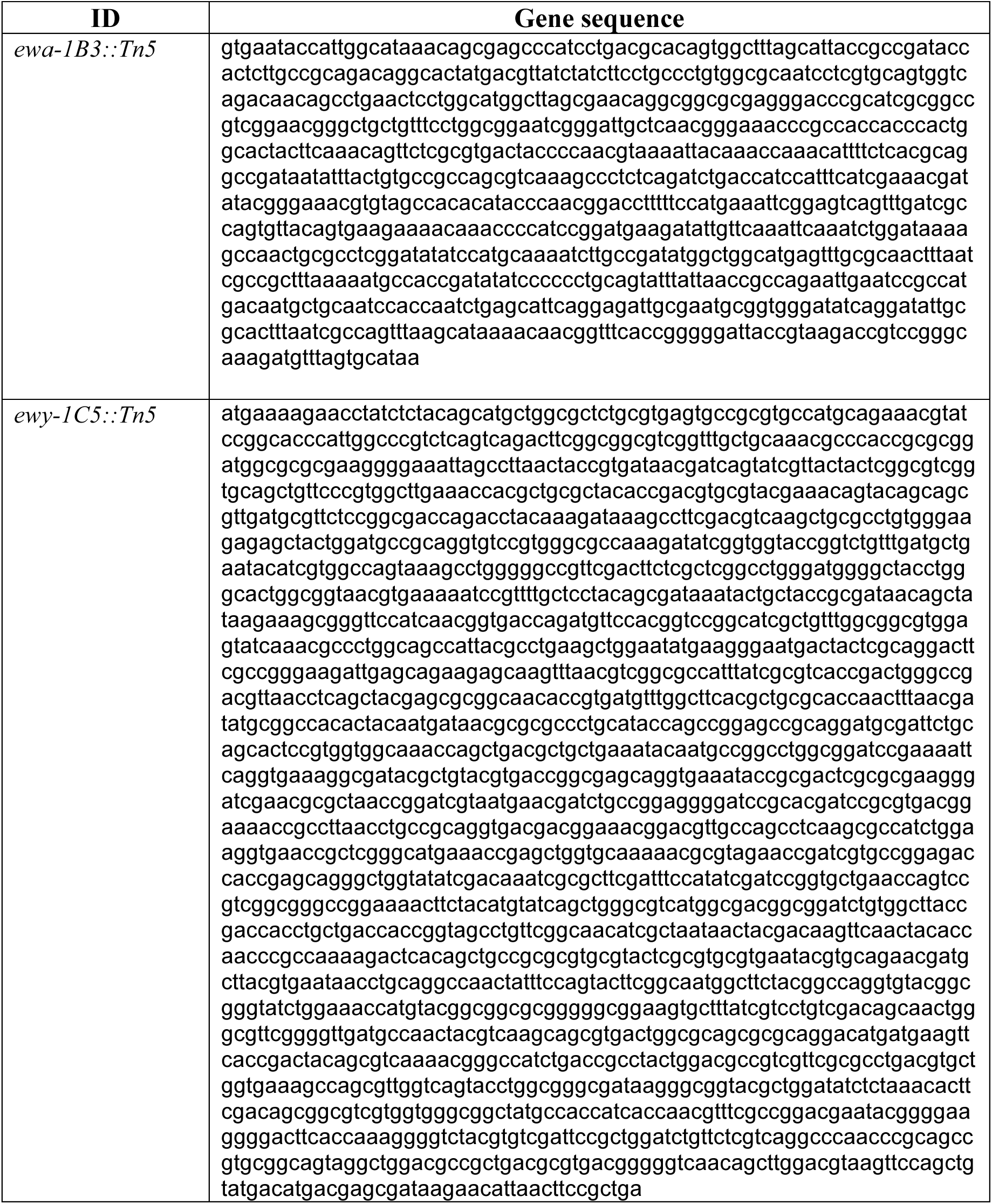
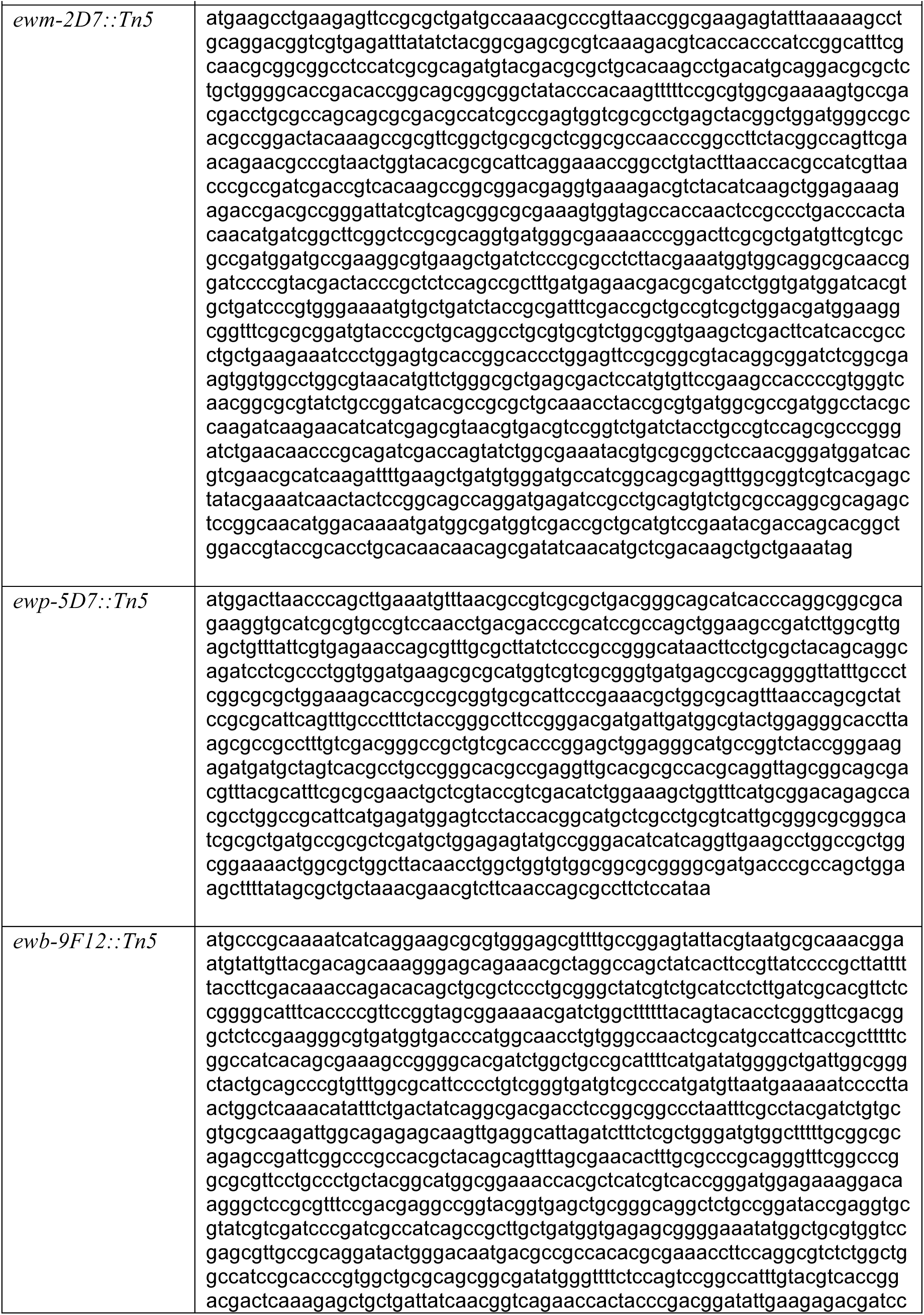
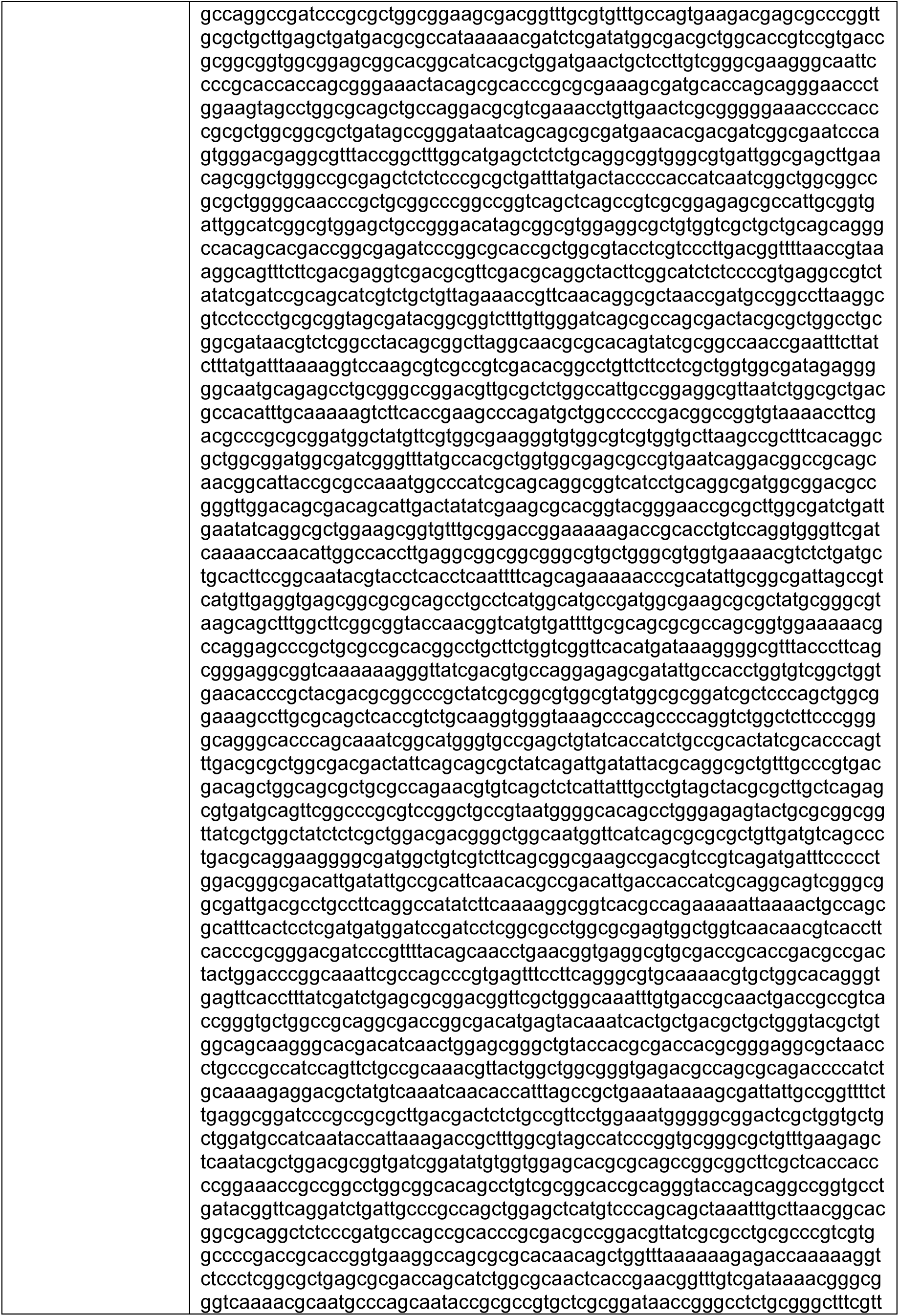
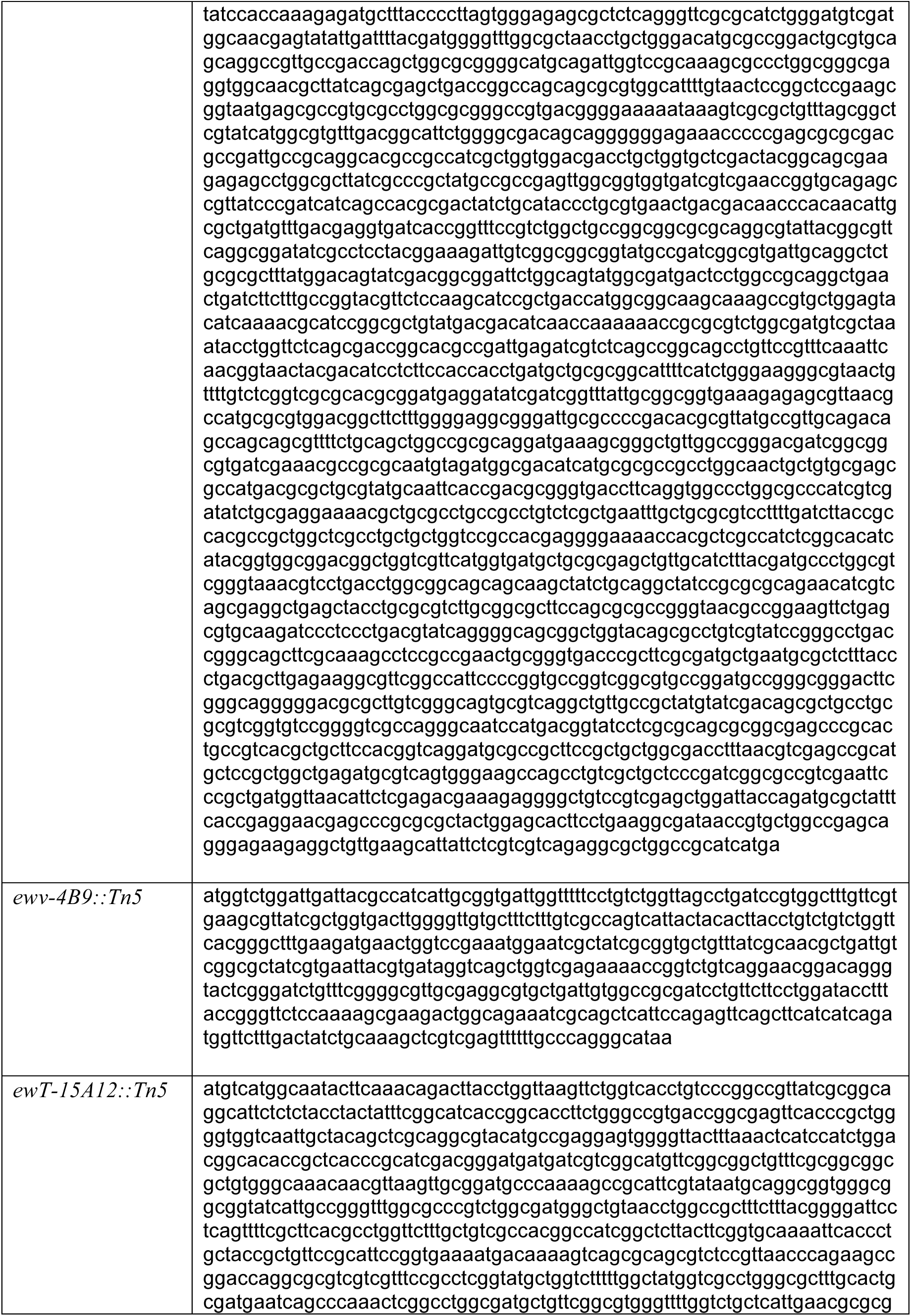
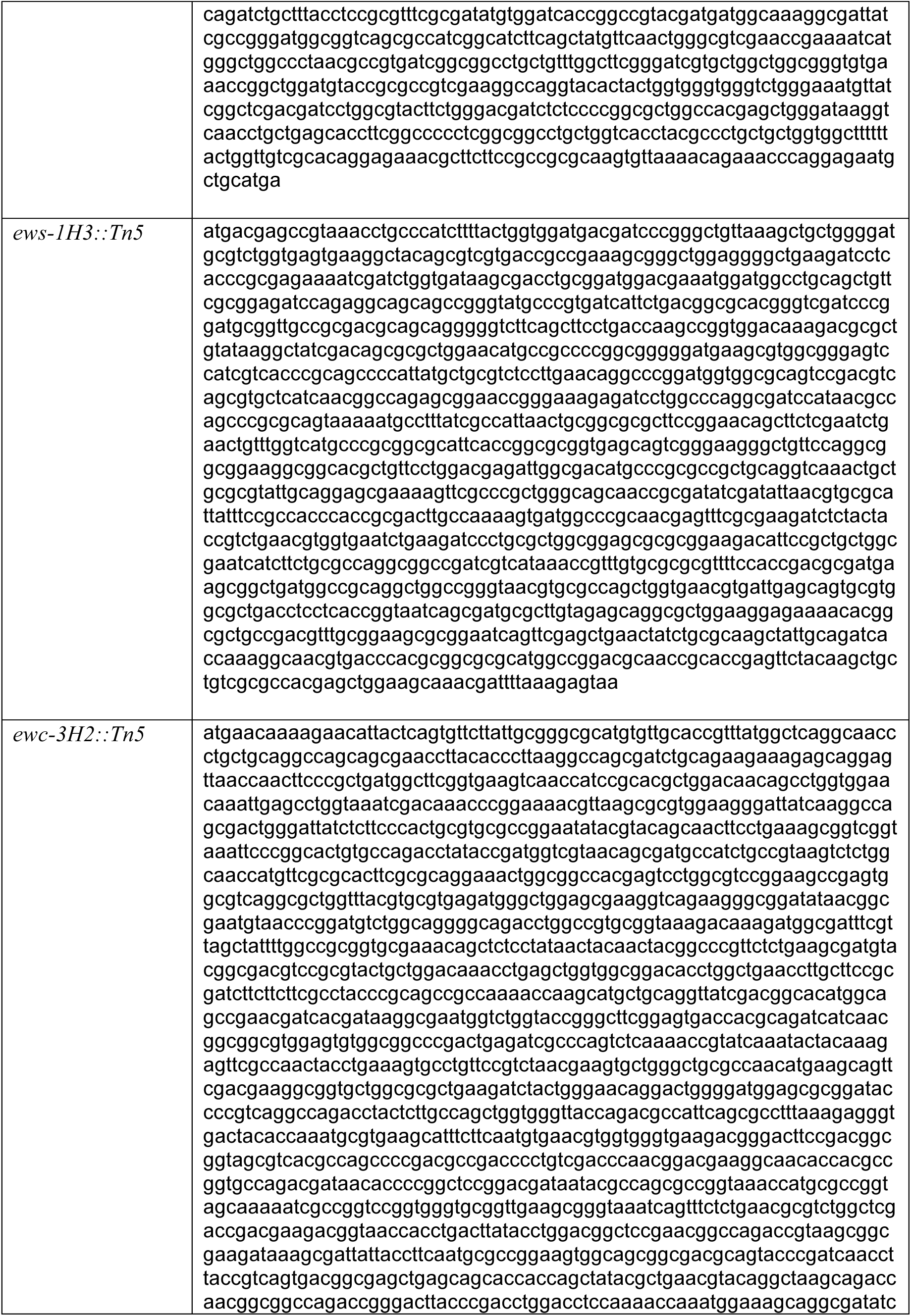
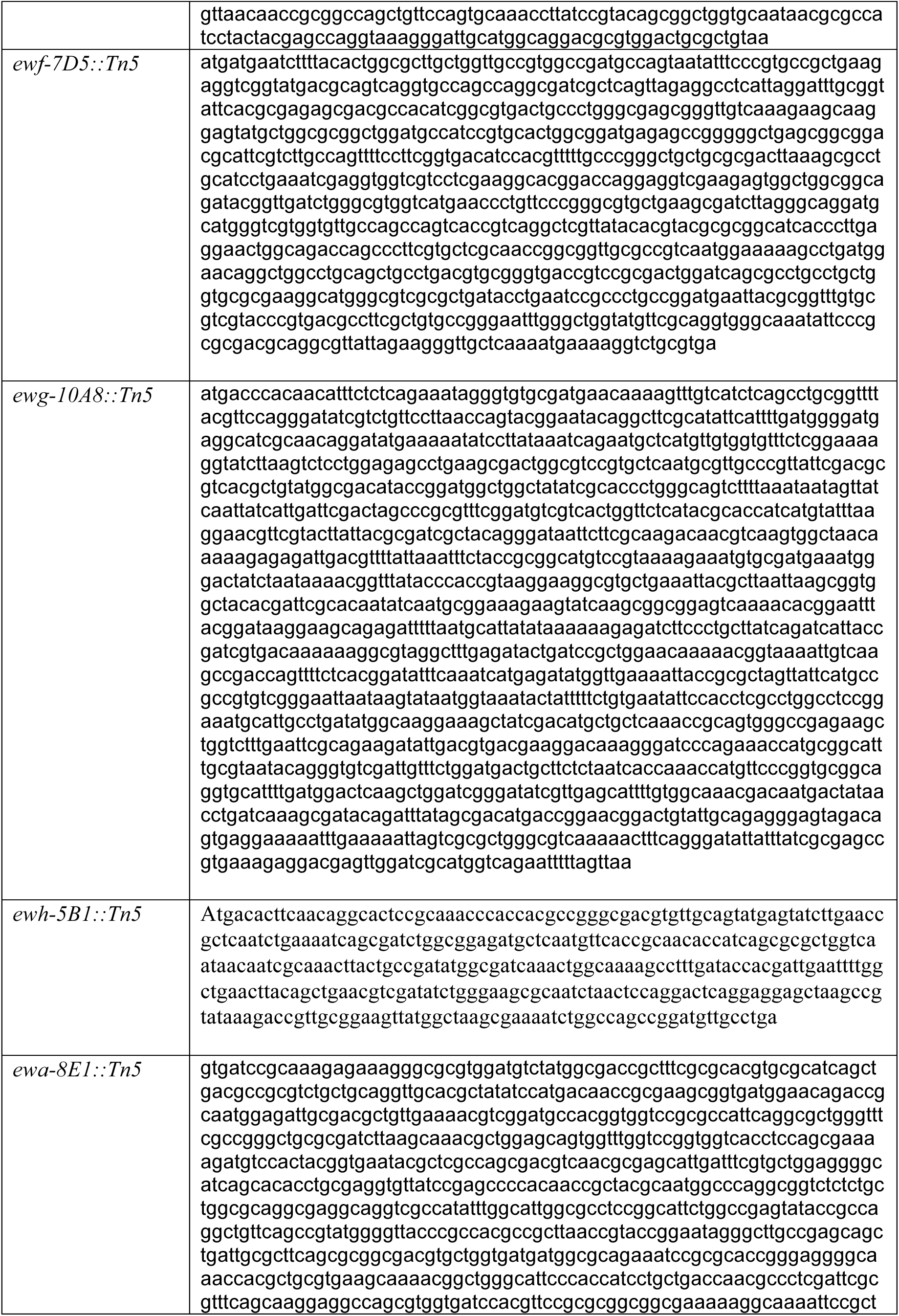
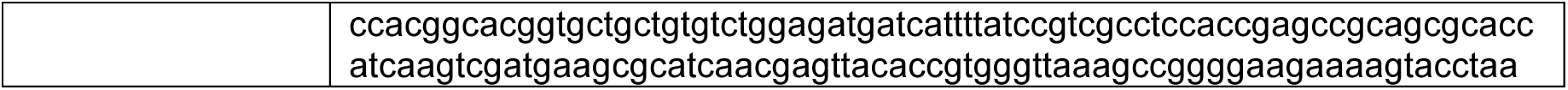
M6 wild type nucleotide coding sequences ^26^ (Ettinger et al, 2015) corresponding to Tn5 insertion mutants that showed loss of antifungal activity against *F. graminearum in vitro* ID Gene sequence

**Supplementary Table 7.**
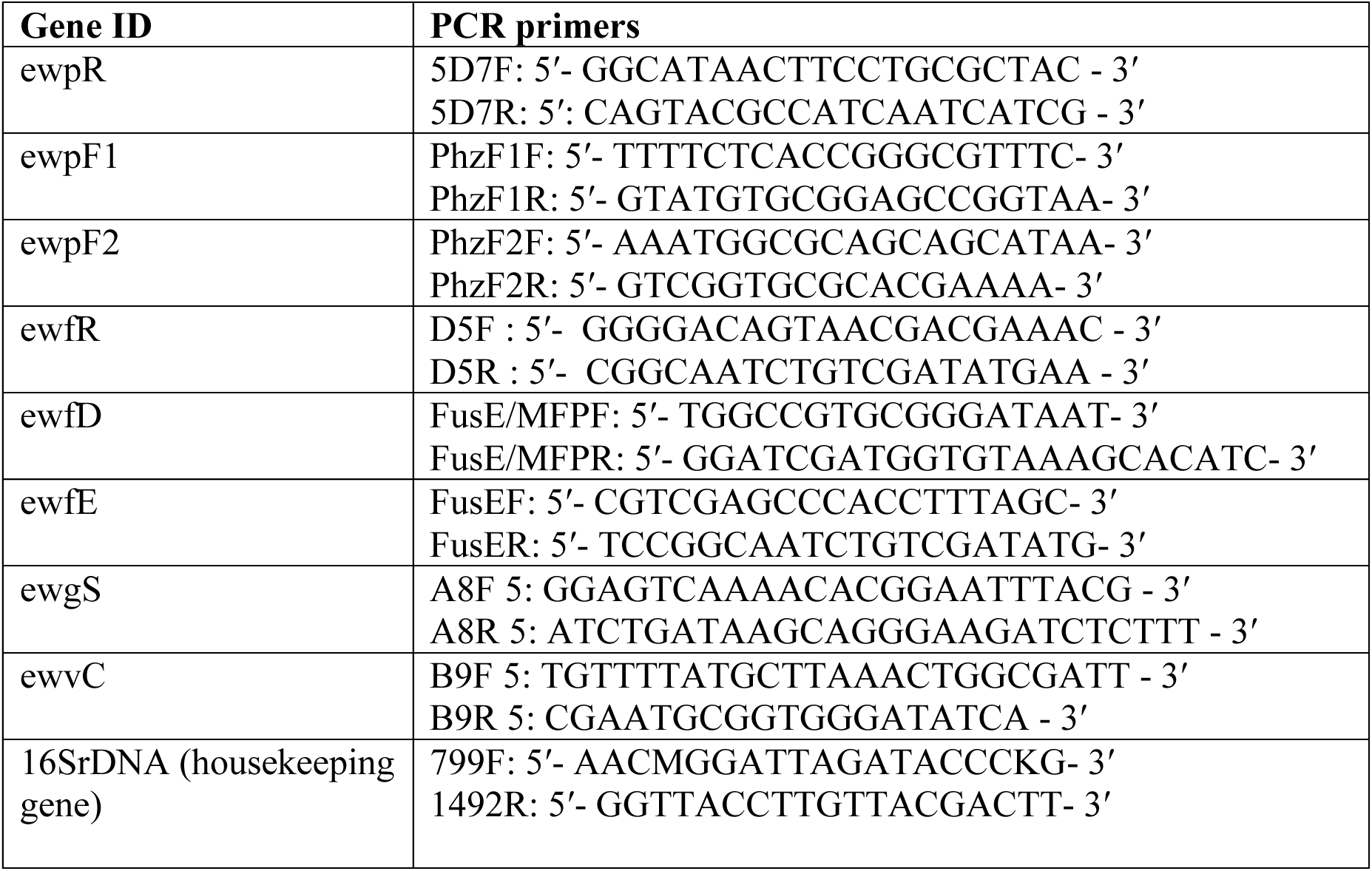
Gene-specific primers used in quantitative real-time PCR analysis.

**Supplementary Figure 1.**
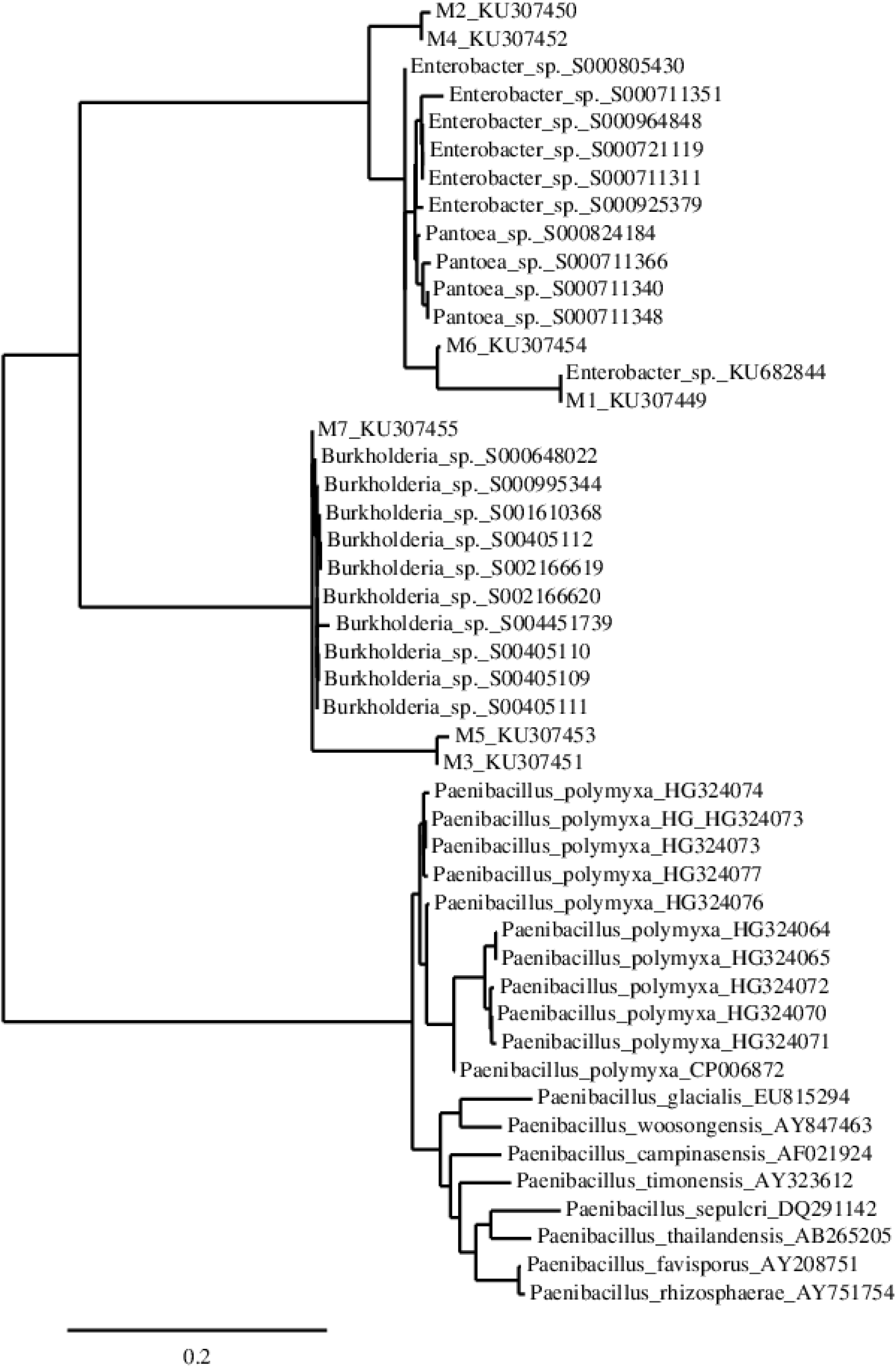
16S rDNA based phylogenetic tree of finger millet endophytes M1-M7.

**Supplementary Figure 2.**
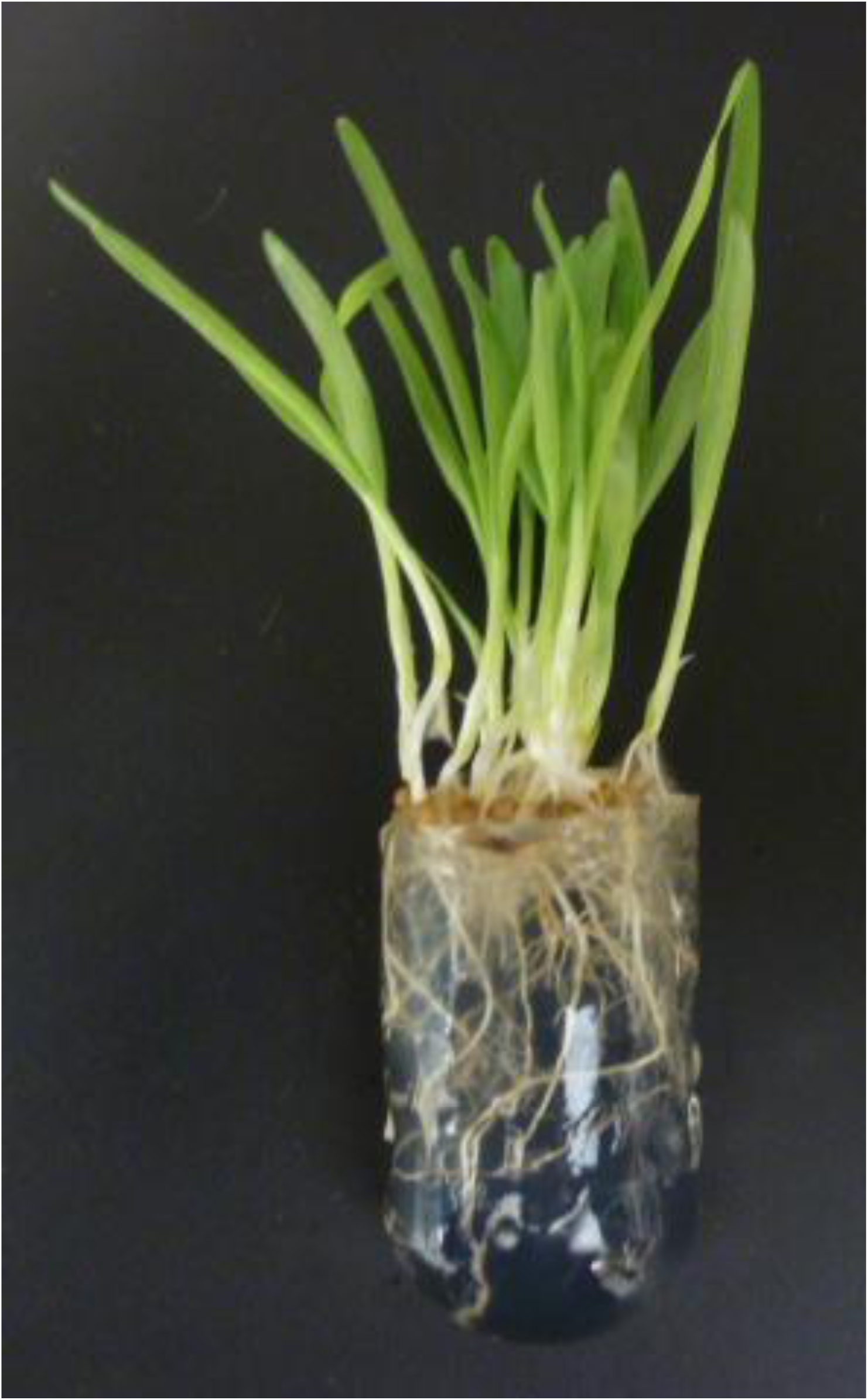
Image of finger millet seedlings previously seed-coated with GFP-tagged M6 showing no pathogenic symptoms, consistent with the strain behaving as an endophyte.

**Supplementary Figure 3.**
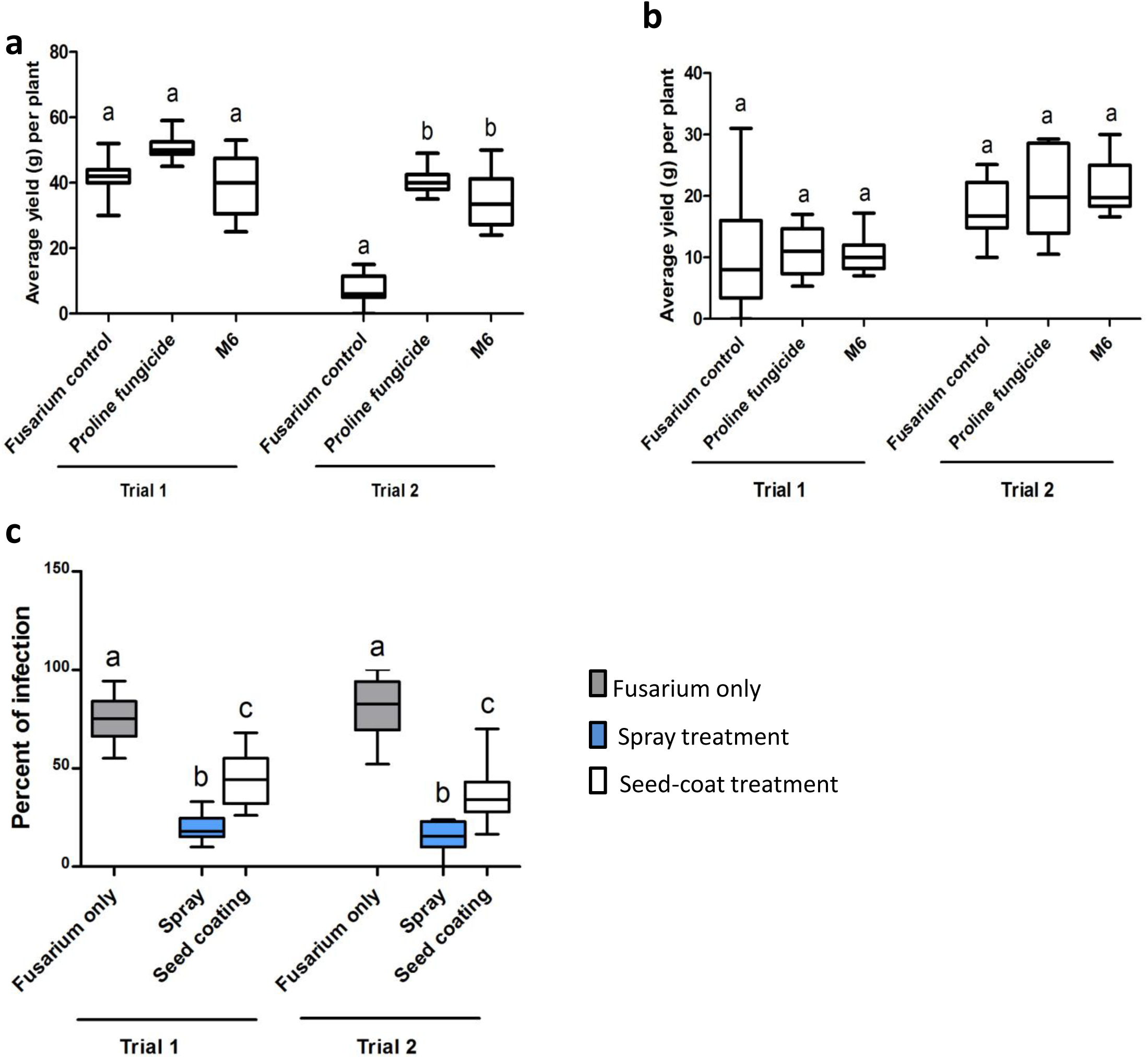
Suppression of *F. graminearum* (Fg) by M6 and its effect on grain yield in greenhouse trials. **a-b**, Effect of endophyte M6 on grain yield per plant in two greenhouse trials for (**a**) maize and (**b**) wheat. **c**, Effect of treatment with endophyte M6 on Fg disease symptoms in maize ears when the endophyte was applied as a seed coat or foliar spray compared to a *Fusarium* only control treatment. Letters that are different from one another within a trial indicate that their means are statistically different (P≤0.05). The whiskers indicate the range of data.

**Supplementary Figure 4.**
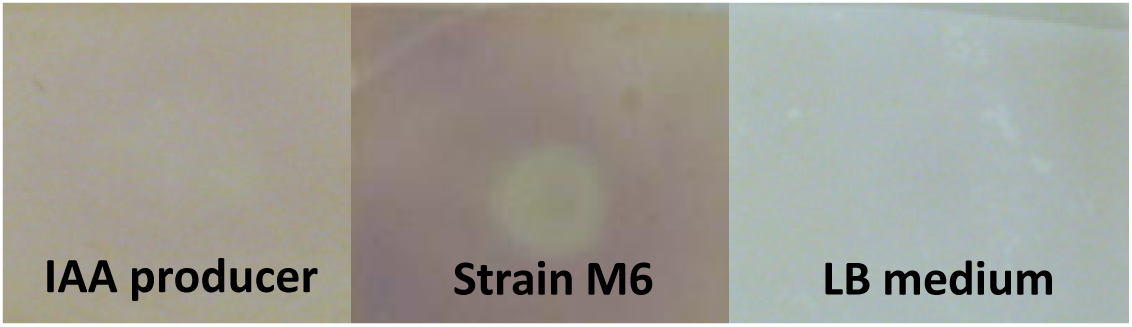
Assay for production of indole-3-acetic acid (IAA, auxin) by wild type strain M6. Production of indole-3-acetic acid (IAA) *in vitro* by wild type strain M6 compared to a positive control (bacterial endophyte strain E10) ^107^ using the Salkowski reagent colorimetric assay.

**Supplementary Figure 5.**
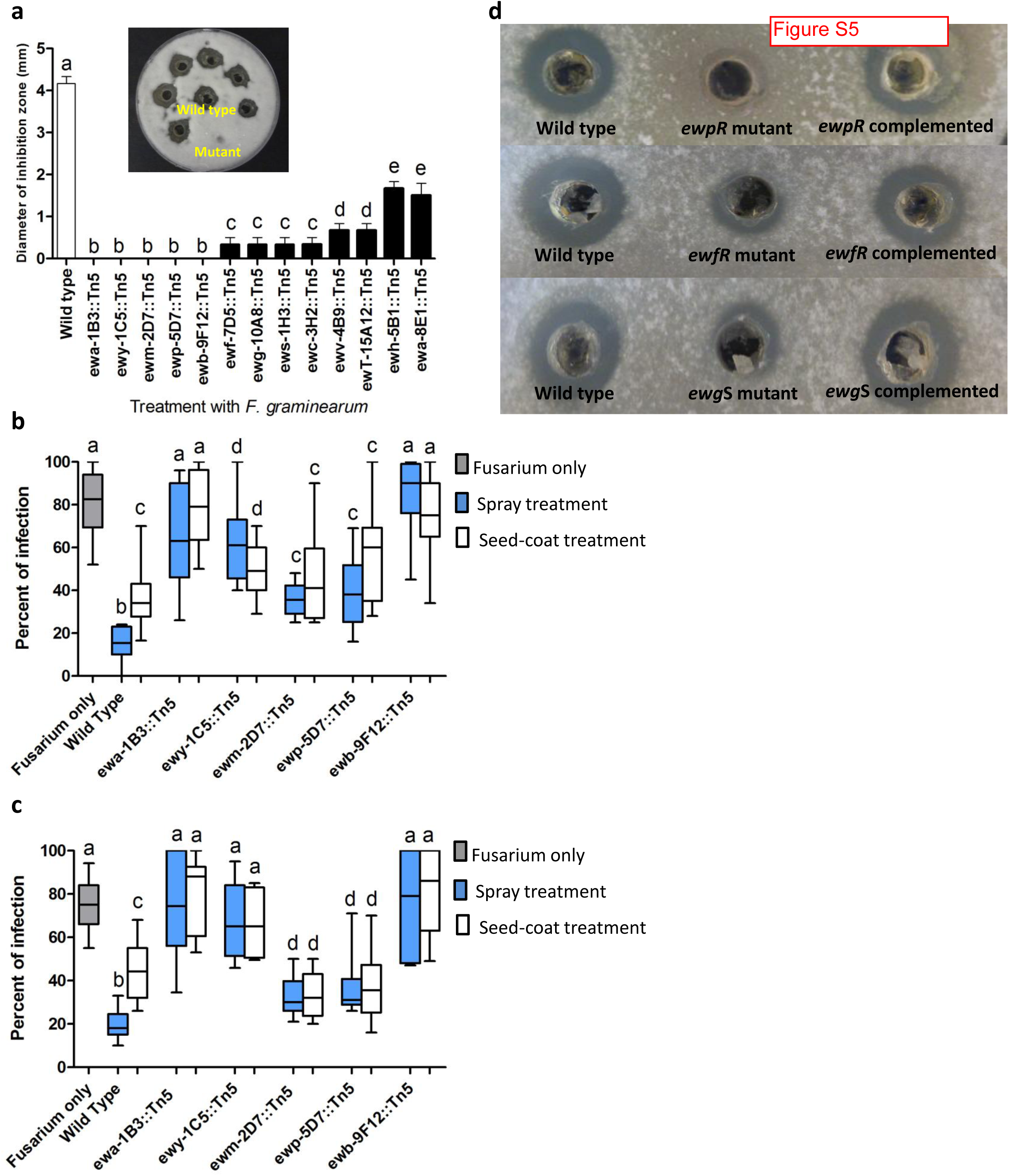
Tn5 mutagenesis-mediated discovery, validation and complementation of genes required for the anti-*Fusarium* activity of strain M6. **a**, Loss of anti-*F.graminearum* activity associated with each Tn5 insertion mutant using the *in vitro* dual culture diffusion assay, along with a representative image (inset) of the mutant screen. **b-c**, *In planta* validation of loss of anti-fungal activity of M6 mutant strains based on quantification of *F. graminearum* disease symptoms on maize ears, in (**b**) greenhouse trial 1, and (**c**) greenhouse trial 2. Only mutant strains that completely lost anti-fungal activity *in vitro* were selected for *in planta* validation. The whiskers indicate the range of data points. Letters that are different from one another indicate that their means are statistically different (P≤0.05). **d**, Genetic complementation of Tn5 mutants with the predicted, corresponding wild type coding sequences. Shown are representative images.

**Supplementary Figure 6.**
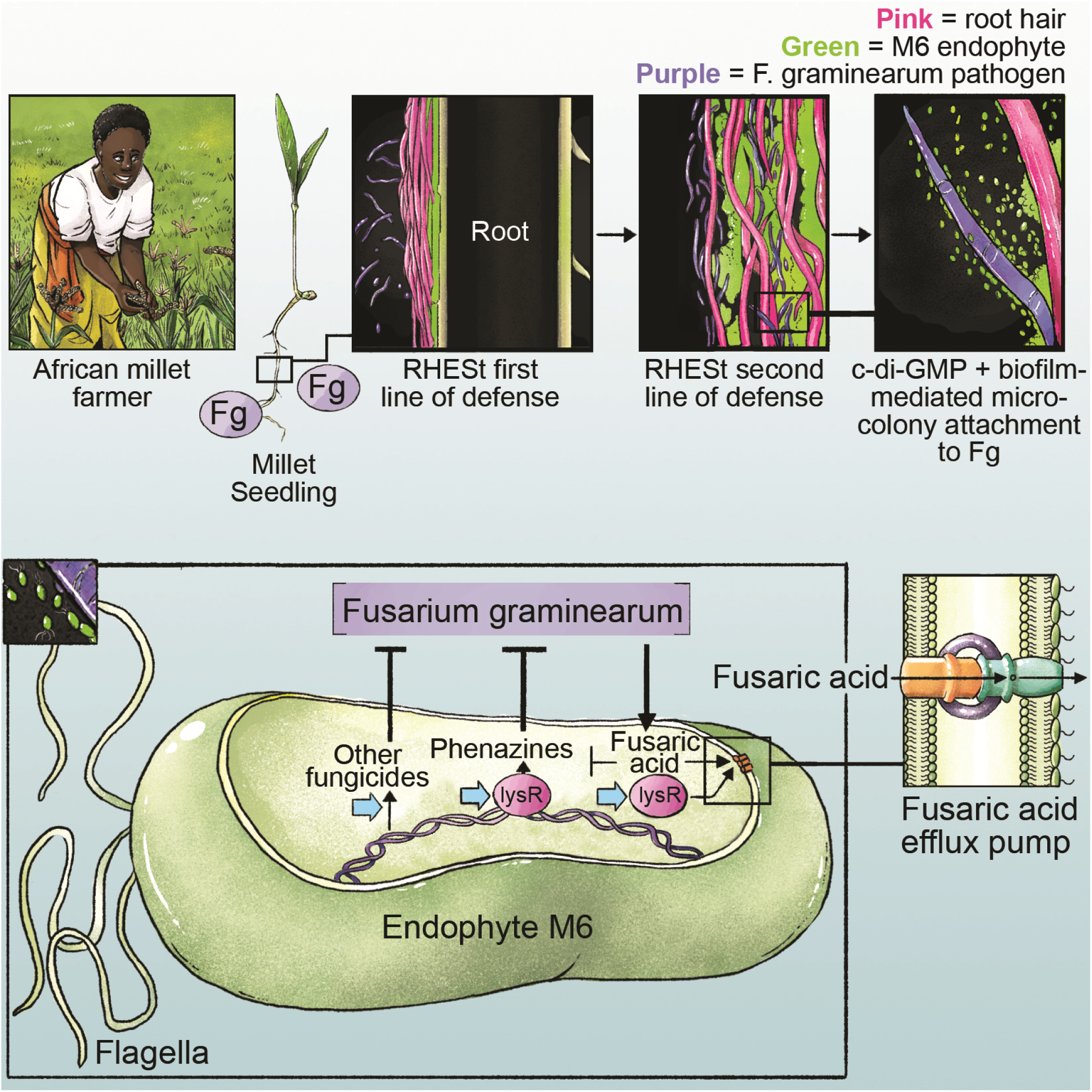
Model to illustrate the interaction between strain M6, the host plant and *F. graminearum* pathogen. Following pathogen sensing, M6 swarm towards *Fusarium* hyphae and induces local hair growth, perhaps mediated by M6-IAA production. M6 then forms microcolony stacks between the elongated and bent root hairs resulting root hair-endophyte stack (RHESt) formation, likely mediated by biofilms. The RHEST formation prevents entry and/or traps *F. graminearum* for subsequent killing. M6 killing requires diverse antimicrobial compounds including phenazines. *Fusarium* will produce fusaric acid which interferes with phenazine biosynthesis. However, M6 has a specialized fusaric acid-resistance pump system which is predicted to pump the mycotoxin outside the endophyte cell.

**Supplementary Figure 7.**
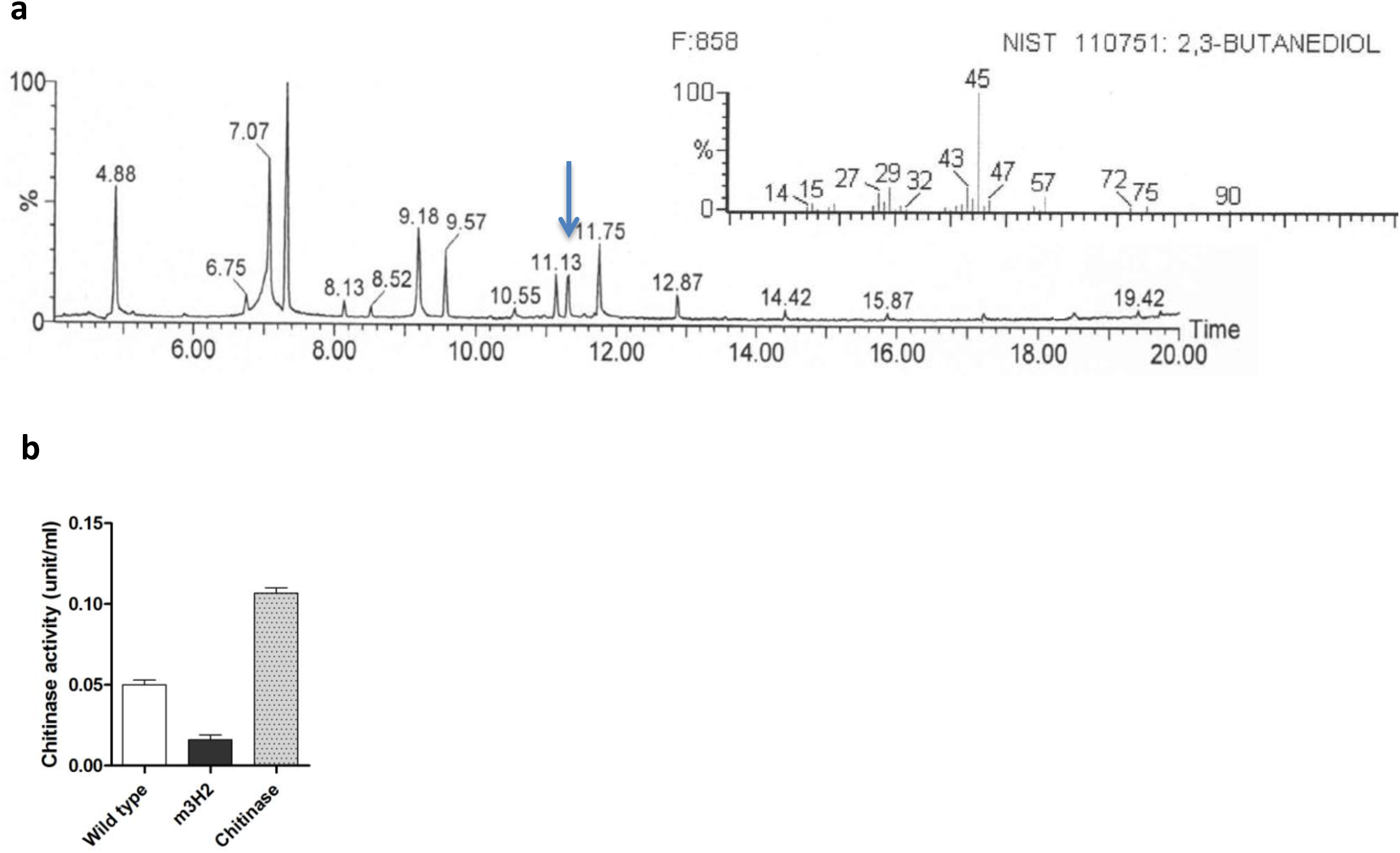
Assays for production of butanediol and chitinase by strain M6. **a**, Entire GC chromatogram showing an arrow pointing to a peak with RT 11.13 with a molecular weight and fragmentation pattern (inset) that matches 2, 3 butanediol when searched against the NIST 2008 database. **b**, Quantification of chitinase production by an M6 mutant strain carrying a Tn5 insertion in a chitinase encoding gene (*ewc-3H2*::Tn5) compared to wild type M6 (see Supplementary Table 7).

## Author Contributions

WKM designed and conducted all experiments, analyzed all data and wrote the manuscript. CS assisted in the greenhouse trials. VLR performed the DON quantification experiments. CE and JE sequenced the M6 genome and provided gene annotations. MNR helped to design the experiments and edited the manuscript. All authors read and approved the manuscript.

## Acknowledgements

We are very thankful to Dr. Michaela Streuder (Department of Molecular and Cellular Biology, University of Guelph) for assistance in confocal microscopy experiments and for her insightful comments. We thank Prof. A. Schaafsma and Dr. Ljiljana Tamburic-Ilincic (Ridgetown College, University of Guelph, Canada) for kindly providing hybrid maize and wheat seeds, respectively. We thank Marina Atalla for assistance in disease scoring. WKM was supported by generous scholarships from the Government of Egypt and the University of Guelph (International Graduate Student Scholarships, 2012, 2014). This research was supported by grants to MNR by the Ontario Ministry of Agriculture, Food and Rural Affairs (OMAFRA), Grain Farmers of Ontario (GFO), Natural Sciences and Engineering Research Council of Canada (NSERC) and the CIFSRF program funded by the International Development Research Centre (IDRC, Ottawa) and Canadian Department of Global Affairs.

## References

1. Goron, T. L. & Raizada, M. N. Genetic diversity and genomic resources available for the small millet crops to accelerate a New Green Revolution. Front. Plant Sci. 6, 157 (2015).

2. Thilakarathna, M. S. & Raizada, M. N. A review of nutrient management studies involving finger millet in the semi-arid tropics of Asia and Africa. Agronomy 5, 262-290 (2015).

3. Hilu, K. W. & de Wet, J. M. J. Domestication of *Eleusine coracana*. Econ. Bot. 30, 199-208 (1976).

4. Mousa, W. K. et al. An endophytic fungus isolated from finger millet (*Eleusine coracana*) produces anti-fungal natural products. Front. Microbiol. 6, 1157(2015).

5. Sundaramari, P. V. a. M. Rationality and adoption of indigenous cultivation practices of finger millet (*Eleusine coracana* (L.) Gaertn.) by the tribal farmers of Tamil Nadu. Intl. J. Manag. Social Sci. 02, 970-977 (2015).

6. Sutton, J. C. Epidemiology of wheat head blight and maize ear rot caused by *Fusarium graminearum*. Can. J. Plant Pathol. 4, 195-209 (1982).

7. Mousa, W. K., Shearer, C., Limay-Rios, V., Zhou, T. & Raizada, M. N. Bacterial endophytes from wild maize suppress *Fusarium graminearum* in modern maize and inhibit mycotoxin accumulation. Front. Plant Sci. 6, 805 (2015).

8. Munimbazi, C. & Bullerman, L. B. Molds and mycotoxins in foods from Burundi. J. Food Prot. 59, 869-875 (1996).

9. Chandrashekar, A. & Satyanarayana, K. Disease and pest resistance in grains of sorghum and millets. J. Cereal Sci. 44, 287-304 (2006).

10. Siwela, M., Taylor, J., de Milliano, W. A. & Duodu, K. G. Influence of phenolics in finger millet on grain and malt fungal load, and malt quality. Food Chem. 121, 443-449 (2010).

11. Wilson, D. Endophyte: the evolution of a term, and clarification of its use and definition. Oikos 73, 274-276 (1995).

12. Johnston-Monje, D. & Raizada, M. N. Conservation and diversity of seed associated endophytes in across boundaries of evolution, ethnography and ecology. PLoS ONE 6, e20396, (2011).

13. Waller, F. et al. The endophytic fungus *Piriformospora indica reprograms barley to salt-stress tolerance, disease resistance, and higher yield*. Proc. NatL. Acad. Sci. (USA) 102, 13386-13391, (2005).

14. Haas, D. & Defago, G. Biological control of soil-borne pathogens by fluorescent pseudomonads. Nat. Rev. Microbiol. 3, 307-319 (2005).

15. Mousa, W. K. & Raizada, M. N. The diversity of anti-microbial secondary metabolites produced by fungal endophytes: An interdisciplinary perspective. Front. Microbiol. 4,65 (2013).

16. Mousa, W. K. & Raizada, M. N. Biodiversity of genes encoding anti-microbial traits within plant associated microbes. Front. Plant Sci. 6, 231(2015).

17. O’Donnell, K. et al. Phylogenetic analyses of RPB1 and RPB2 support a middle Cretaceous origin for a clade comprising all agriculturally and medically important fusaria. Fungal Genet. Biol. 52, 20-31 (2013).

18. Saleh, A. A., Esele, J., Logrieco, A., Ritieni, A. & Leslie, J. F. *Fusarium verticillioide*s from finger millet in Uganda. Food Addit. Contam. 29, 1762-1769 (2012).

19. Pall, B. & Lakhani, J. Seed mycoflora of ragi, Eleusine coracana (L.) Gaertn. Res. Develop. Rep. 8, 78-79 (1991).

20. Amata, R. et al. An emended description of *Fusarium brevicatenulatum* and *F. pseudoanthophilum* based on isolates recovered from millet in Kenya. Fungal Divers. 43, 11-25 (2010).

21. Penugonda, S., Girisham, S. & Reddy, S. Elaboration of mycotoxins by seed-borne fungi of finger millet (*Eleusine coracana* L.). Intl. J. Biotech. Mol. Biol. Res. 1, 62-64 (2010).

22. Ramana, M. V., Nayaka, S. C., Balakrishna, K., Murali, H. & Batra, H. A novel PCR–DNA probe for the detection of fumonisin-producing *Fusarium* species from major food crops grown in southern India. Mycology 3, 167-174 (2012).

23. Adipala, E. Seed-borne fungi of finger millet. E. Afr. Agricult. Forest. J. 57, 173-176 (1992).

24. Dereeper, A. et al. Phylogeny. fr: robust phylogenetic analysis for the non-specialist. Nucleic acids Res. 36, 465-469 (2008).

25. Dereeper, A., Audic, S., Claverie, J.-M. & Blanc, G. BLAST-explorer helps you building datasets for phylogenetic analysis. BMC Evol. Biol. 10, 8 (2010).

26. Ettinger, C. L., Mousa, W. M., Raizada, M. N. & Eisen, J. A. Draft Genome Sequence of *Enterobacter* sp. Strain UCD-UG_FMILLET (Phylum Proteobacteria). Genome Announc. 3,e0146-14 (2015).

27. Chongo, G. et al. Reaction of seedling roots of 14 crop species to *Fusarium graminearum* from wheat heads. Can. J. Plant Pathol. 23, 132-137 (2001).

28. Pitts, R. J., Cernac, A. & Estelle, M. Auxin and ethylene promote root hair elongation in Arabidopsis. Plant J. 16, 553-560 (1998).

29. Parsons, J. F. et al. Structure and function of the phenazine biosynthesis protein PhzF from *Pseudomonas fluorescens* 2-79. Biochemistry 43, 12427-12435 (2004).

30. Lu, J. et al. LysR family transcriptional regulator PqsR as repressor of pyoluteorin biosynthesis and activator of phenazine-1-carboxylic acid biosynthesis in *Pseudomonas* sp. M18. J. Biotechnol. 143, 1-9 (2009).

31. 52. Goethals, K., Van Montagu, M. & Holsters, M. Conserved motifs in a divergent nod box of Azorhizobium caulinodans ORS571 reveal a common structure in promoters regulated by LysR-type proteins. Proc. Natl. Acad. Sci. (USA) 89, 1646-1650 (1992).

32. Wang, Y., Luo, Q., Zhang, X. & Wang, W. Isolation and purification of a modified phenazine, griseoluteic acid, produced by *Streptomyces griseoluteus* P510. Res. Microbiol. 162, 311-319, (2011).

33. Nakamura, S., Wang, E. L., Murase, M., Maeda, K. & Umezawa, H. Structure of griseolutein A. J. Antibiot. 12, 55-58 (1959).

34. Giddens, S. R., Feng, Y. & Mahanty, H. K. Characterization of a novel phenazine antibiotic gene cluster in *Erwinia herbicola* Eh1087. Mol. Microbiol. 45, 769-783 (2002).

35. Kitahara, M. et al. Saphenamycin, a novel antibiotic from a strain of Streptomyces. J. Antibiot. 35, 1412-1414 (1982).

36. Toyoda, H., Katsuragi, K., Tamai, T. & Ouchi, S. DNA Sequence of genes for detoxification of fusaric acid, a wilt-inducing agent produced by *Fusarium* species. J. Phytopathol. 133, 265-277 (1991).

37. Utsumi, R. et al. Molecular cloning and characterization of the fusaric acid-resistance gene from *Pseudomonas cepacia*. *Agri*. Biol. Chem. 55, 1913-1918 (1991).

38. Marre, M., Vergani, P. & Albergoni, F. Relationship between fusaric acid uptake and its binding to cell structures by leaves of *Egeria densa* and its toxic effects on membrane permeability and respiration. Physiol. Mol. Plant Pathol. 42, 141-157 (1993).

39. Bacon, C., Hinton, D. & Hinton, A. Growth-inhibiting effects of concentrations of fusaric acid on the growth of *Bacillus mojavensis* and other biocontrol *Bacillus* species. J. App. Microbiol. 100, 185-194 (2006).

40. Borges-Walmsley, M., McKeegan, K. & Walmsley, A. Structure and function of efflux pumps that confer resistance to drugs. Biochem. J. 376, 313-338 (2003).

41. Hu, R.-M., Liao, S.-T., Huang, C.-C., Huang, Y.-W. & Yang, T.-C. An inducible fusaric acid tripartite efflux pump contributes to the fusaric acid resistance in *Stenotrophomonas maltophilia*. PLoS ONE 7, e51053 (2012).

42. van Rij, E. T., Girard, G., Lugtenberg, B. J. & Bloemberg, G. V. Influence of fusaric acid on phenazine-1-carboxamide synthesis and gene expression of *Pseudomonas chlororaphis* strain PCL1391. Microbiology 151, 2805-2814 (2005).

43. Römling, U., Galperin, M. Y. & Gomelsky, M. Cyclic di-GMP: the first 25 years of a universal bacterial second messenger. Microbiol. Mol. Biol. Rev.77, 1-52 (2013).

44. Fath, M., Mahanty, H. & Kolter, R. Characterization of a purF operon mutation which affects colicin V production. J. Bacteriol. 171, 3158-3161 (1989).

45. Fath, M. J., Zhang, L. H., Rush, J. & Kolter, R. Purification and characterization of colicin V from *Escherichia coli* culture supernatants. Biochemistry 33, 6911-6917 (1994).

46. Soliman, S. S. et al. An endophyte constructs fungicide-containing extracellular barriers for its host plant. Curr. Biol. 25, 2570-2576 (2015).

47. Reinhold-Hurek, B., Bünger, W., Burbano, C. S., Sabale, M. & Hurek, T. Roots shaping their microbiome: global hotspots for microbial activity. Ann. Rev. Phytopathol. 53, 403-424 (2015).

48. Compant, S., Clément, C. & Sessitsch, A. Plant growth-promoting bacteria in the rhizo-and endosphere of plants: their role, colonization, mechanisms involved and prospects for utilization. Soil Biol. Biochem. 42, 669-678 (2010).

49. Glaeser, S. P. et al. Non-pathogenic *Rhizobium radiobacter* F4 deploys plant beneficial activity independent of its host *Piriformospora indica*. ISME J. in press (2015).

50. Germaine, K. J., Keogh, E., Ryan, D. & Dowling, D. N. Bacterial endophyte-mediated naphthalene phytoprotection and phytoremediation. FEMS Microbiol. Lett. 296, 226-234 (2009).

51. Marasco, R. et al. A drought resistance-promoting microbiome is selected by root system under desert farming. PloS ONE 7, e48479 (2012).

52. Dandurand, L., Schotzko, D. & Knudsen, G. Spatial patterns of rhizoplane populations of *Pseudomonas fluorescens*. Appl. Environmen. Microbiol. 63, 3211-3217 (1997).

53. Foster, R. The ultrastructure of the rhizoplane and rhizosphere. Ann. Rev. Phytopathol. 24, 211-234 (1986).

54. Fukui, R. et al. Spatial colonization patterns and interaction of bacteria on inoculated sugar beet seed. Phytopathology 84, 1338-1345 (1994).

55. Rovira, A. A study of the development of the root surface microflora during the initial stages of plant growth. J. Appl. Bacteriol. 19, 72-79 (1956).

56. Rovira, A. & Campbell, R. Scanning electron microscopy of microorganisms on the roots of wheat. Microb. Ecol. 1, 15-23 (1974).

57. Newman, E. & Bowen, H. Patterns of distribution of bacteria on root surfaces. Soil Biol. Biochem. 6, 205-209 (1974).

58. Mercado-Blanco, J. & Prieto, P. Bacterial endophytes and root hairs. Plant Soil 361, 301-306 (2012).

59. Danhorn, T. & Fuqua, C. Biofilm formation by plant-associated bacteria. Annu. Rev. Microbiol. 61, 401-422 (2007).

60. Lee, R. D.-W. & Cho, H.-T. Auxin, the organizer of the hormonal/environmental signals for root hair growth. Front. Plant Sci. 4, 448 (2013).

61. Patten, C. L. & Glick, B. R. Bacterial biosynthesis of indole-3-acetic acid. Can. J. Microbiol. 42, 207-220 (1996).

62. Oldroyd, G. E. Speak, friend, and enter: signalling systems that promote beneficial symbiotic associations in plants. Nat. Rev. Microbiol. 11, 252-263 (2013).

63. Du, X., Li, Y., Zhou, Q. & Xu, Y. Regulation of gene expression in *Pseudomonas aeruginosa* M18 by phenazine-1-carboxylic acid. Appl. Microbiol. Biotechnol. 99, 813-825 (2015).

64. Srivastava, D. & Waters, C. M. A tangled web: regulatory connections between quorum sensing and cyclic di-GMP. J. Bacteriol. 194, 4485-4493 (2012).

65. Flemming, H.-C. & Wingender, J. The biofilm matrix. Nat. Rev. Microbiol. 8, 623-633 (2010).

66. An, S.Q. et al. Novel cyclic di-GMP effectors of the YajQ protein family control bacterial virulence. PLoS Pathog. 10, e1004429 (2014).

67. Landini, P., Antoniani, D., Burgess, J. G. & Nijland, R. Molecular mechanisms of compounds affecting bacterial biofilm formation and dispersal. Appl. Microbiol. Biotechnol. 86, 813-823 (2010).

68. Steinberg, N. & Kolodkin-Gal, I. The matrix reloaded: probing the extracellular matrix synchronizes bacterial communities. J. Bacteriol. 197, 2092–2103 (2015).

69. Cortes-Barco, A., Hsiang, T. & Goodwin, P. Induced systemic resistance against three foliar diseases of *Agrostis stolonifera* by (2R, 3R)-butanediol or an isoparaffin mixture. Annal. Appl. Biol. 157, 179-189 (2010).

70. Mavrodi, D. V. et al. Recent insights into the diversity, frequency and ecological roles of phenazines in fluorescent *Pseudomonas* spp. Environmen. Microbiol. 15, 675-686 (2013).

71. Gurusiddaiah, S., Weller, D., Sarkar, A. & Cook, R. Characterization of an antibiotic produced by a strain of *Pseudomonas fluorescens* inhibitory to *Gaeumannomyces graminis* var. tritici and *Pythium* spp. Antimicrob. Agents Chemotherap.29, 488-495 (1986).

72. Anjaiah, V. et al. Involvement of phenazines and anthranilate in the antagonism of *Pseudomonas aeruginosa* PNA1 and Tn5 Derivatives toward *Fusarium* spp. and *Pythium* spp. Mol. Plant-Microb. Interact. 11, 847-854 (1998).

73. Bloemberg, G. V. & Lugtenberg, B. J. Phenazines and their role in biocontrol by Pseudomonas bacteria. New Phytol. 157, 503-523 (2003).

74. Hori, M., Nozaki, S., Nagami, K., Asakura, H. & Umezawa, H. Inhibition of DNA synthesis by griseolutein in *Escherichia coli* through a possible interaction at the cell surface. Biochimica et Biophysica Acta (BBA)-Nuc. Acids Protein Synth. 521, 101-110 (1978).

75. Hassan, H. M. & Fridovich, I. Mechanism of the antibiotic action pyocyanine. J. Bacteriol. 141, 156-163 (1980).

76. Hassett, D. J., Schweizer, H. P. & Ohman, D. E. *Pseudomonas aeruginosa* sodA and sodB mutants defective in manganese-and iron-cofactored superoxide dismutase activity demonstrate the importance of the iron-cofactored form in aerobic metabolism. J. Bacteriol. 177, 6330-6337 (1995).

77. Audenaert, K., Pattery, T., Cornelis, P. & Hofte, M. Induction of systemic resistance to *Botrytis cinerea* in tomato by *Pseudomonas aeruginosa* 7NSK2: role of salicylic acid, pyochelin, and pyocyanin. Mol. Plant. Microbe Interact. 15, 1147-1156 (2002).

78. Price-Whelan, A., Dietrich, L. E. & Newman, D. K. Rethinking 'secondary'metabolism: physiological roles for phenazine antibiotics. Nat. Chem. Biol. 2, 71-78 (2006).

79. Wang, Y. & Newman, D. K. Redox reactions of phenazine antibiotics with ferric (hydr) oxides and molecular oxygen. Environ. Sci. Technol. 42, 2380-2386 (2008).

80. Hernandez, M. & Newman, D. Extracellular electron transfer. Cell. Mol. Life Sci. 58, 1562-1571 (2001).

81. Fu, Y.-C., Zhang, T. & Bishop, P. Determination of effective oxygen diffusivity in biofilms grown in a completely mixed biodrum reactor. Water Sci. Technol. 29, 455-462 (1994).

82. Stewart, P. S. Diffusion in biofilms. J. Bacteriol. 185, 1485-1491 (2003).

83. Whitchurch, C. B., Tolker-Nielsen, T., Ragas, P. C. & Mattick, J. S. Extracellular DNA required for bacterial biofilm formation. Science 295, 1487-1487 (2002).

84. Petersen, F. C., Tao, L. & Scheie, A. A. DNA binding-uptake system: a link between cell-to-cell communication and biofilm formation. J. Bacteriol. 187, 4392-4400 (2005).

85. Das, T., Sharma, P. K., Busscher, H. J., van der Mei, H. C. & Krom, B. P. Role of extracellular DNA in initial bacterial adhesion and surface aggregation. Appl. Environ. Microbiol. 76, 3405-3408 (2010).

86. Das, T., Kutty, S. K., Kumar, N. & Manefield, M. Pyocyanin facilitates extracellular DNA binding to *Pseudomonas aeruginosa* influencing cell surface properties and aggregation. PloS ONE 8, e58299 (2013).

87. Das, T. & Manefield, M. Pyocyanin promotes extracellular DNA release in *Pseudomonas aeruginosa*. PLoS ONE 7, e46718. (2012).

88. Das, T. et al. Phenazine virulence factor binding to extracellular DNA is important for *Pseudomonas aeruginosa* biofilm formation. Sci. Rep. 5, 8398 (2015).

89. Pierson III, L. S. & Pierson, E. A. Metabolism and function of phenazines in bacteria: impacts on the behavior of bacteria in the environment and biotechnological processes. Appl. Microbiol. Biotechnol. 86, 1659-1670 (2010).

90. Gérard, F., Pradel, N. & Wu, L.-F. Bactericidal activity of colicin V is mediated by an inner membrane protein, SdaC, of *Escherichia coli*. J. Bacteriol. 187, 1945-1950 (2005).

91. Gillor, O. & Ghazaryan, L. Recent advances in bacteriocin application as antimicrobials. Recent Pat. Anti-infect. Drug Discov. 2, 115-122 (2007).

92. Hammami, I., Triki, M. & Rebai, A. Purification and characterization of the novel bacteriocin bac ih7 with antifungal and antibacterial properties. J. Plant Pathol. 93, 443-454 (2011).

93. Dahiya, N., Tewari, R. & Hoondal, G. S. Biotechnological aspects of chitinolytic enzymes: a review. Appl. Microbiol. Biotechnol. 71, 773-782 (2005).

94. Dahiya, N., Tewari, R., Tiwari, R. P. & Hoondal, G. S. Production of an antifungal chitinase from *Enterobacter* sp. NRG4 and its application in protoplast production. World J. Microbiol. Biotechnol. 21, 1611-1616 (2005).

95. Burkhead, K. D., Slininger, P. J. & Schisler, D. A. Biological control bacterium *Enterobacter cloacae* S11:T:07 (NRRL B-21050) produces the antifungal compound phenylacetic acid in Sabouraud maltose broth culture. Soil Biol. Biochem. 30, 665-667 (1998).

96. Slininger, P., Burkhead, K. & Schisler, D. Antifungal and sprout regulatory bioactivities of phenylacetic acid, indole-3-acetic acid, and tyrosol isolated from the potato dry rot suppressive bacterium *Enterobacter cloacae* S11: T: 07. J. Ind. Microbiol. Biotechnol. 31, 517-524 (2004).

97. Schisler, D. A., Slininger, P. J., Kleinkopf, G., Bothast, R. J. & Ostrowski, R. C. Biological control of *Fusarium* dry rot of potato tubers under commercial storage conditions. Am. J. Potato Res. 77, 29-40 (2000).

98. Herbert, R. B. & Knaggs, A. R. Biosynthesis of the antibiotic obafluorin from p-aminophenylalanine and glycine (glyoxylate). J. Chem. Soci. (Perkin Transactions) 1, 109-113 (1992).

99. Lewis, E. A., Adamek, T. L., Vining, L. C. & White, R. L. Metabolites of a blocked chloramphenicol producer. J. Nat. Prod. 66, 62-66 (2003).

100. He, J., Magarvey, N., Piraee, M. & Vining, L. The gene cluster for chloramphenicol biosynthesis in *Streptomyces venezuelae* ISP5230 includes novel shikimate pathway homologues and a monomodular non-ribosomal peptide synthetase gene. Microbiology 147, 2817-2829 (2001).

101. Wang, K., Kang, L., Anand, A., Lazarovits, G. & Mysore, K. S. Monitoring *in planta* bacterial infection at both cellular and whole-plant levels using the green fluorescent protein variant GFPuv. New Phytol. 174, 212-223 (2007).

102. 124. Solovyev, V. & Salamov, A. Automatic annotation of microbial genomes and metagenomic sequences. in Metagenomics and its Applications in Agriculture, Biomedicine and Environmental Studies (ed. Robert, W. Li), 61-78 (Nova Science, 2011).

103. de Jong, A., Pietersma, H., Cordes, M., Kuipers, O. P. & Kok, J. PePPER: a webserver for prediction of prokaryote promoter elements and regulons. BMC Genom. 13, 299 (2012).

104. Stepanović, S., Vuković, D., Dakić, I., Savić, B. & Švabić-Vlahović, M. A modified microtiter-plate test for quantification of staphylococcal biofilm formation. J. Microbiol. Methods 40, 175-179 (2000).

105. Bacon, C., Hinton, D. & Hinton, A. Growth-inhibiting effects of concentrations of fusaric acid on the growth of *Bacillus mojavensis* and other biocontrol *Bacillus* species. J. Appl. Microbiol. 100, 185-194 (2006).

106. Paulus, H. & Gray, E. The biosynthesis of polymyxin B by growing cultures of *Bacillus polymyxa*. J. Biol. Chem. 239, 865-871 (1964).

107. Shehata, H. Molecular and physiological mechanisms underlying the antifungal and nutrient acquisition activities of beneficial microbes. PhD thesis, University of Guelph, ON Canada. (2016).

